# Cholinergic modulation of dopamine release drives effortful behavior

**DOI:** 10.1101/2025.06.18.660394

**Authors:** Gavin C. Touponse, Matthew B. Pomrenze, Teema Yassine, Viraj Mehta, Nicholas Denomme, Zihui Zhang, Robert C. Malenka, Neir Eshel

**Affiliations:** Department of Psychiatry and Behavioral Sciences, Stanford University, Stanford, CA 94305; Nancy Pritzker Laboratory, Department of Psychiatry and Behavioral Sciences, Stanford University, Stanford, CA 94305

## Abstract

Effort is costly: given a choice, we tend to avoid it^1^. But in many cases, effort adds value to the ensuing rewards^2^. From ants^3^ to humans^4^, individuals prefer rewards that had been harder to achieve. This counterintuitive process may promote reward-seeking even in resource-poor environments, thus enhancing evolutionary fitness^5^. Despite its ubiquity, the neural mechanisms supporting this behavioral effect are poorly understood. Here we show that effort amplifies the dopamine response to an otherwise identical reward, and this amplification depends on local modulation of dopamine axons by acetylcholine. High-effort rewards evoke rapid acetylcholine release from local interneurons in the nucleus accumbens. Acetylcholine then binds to nicotinic receptors on dopamine axon terminals to augment dopamine release when reward is delivered. Blocking the cholinergic modulation blunts dopamine release selectively in high-effort contexts, impairing effortful behavior while leaving low-effort reward consumption intact. These results reconcile *in vitro* studies, which have long demonstrated that acetylcholine can trigger dopamine release directly through dopamine axons^6–11;^ with *in vivo* studies that failed to observe such modulation^12–14^, but did not examine high-effort contexts. Our findings uncover a mechanism that drives effortful behavior through context-dependent local interactions between acetylcholine and dopamine axons.

## Main

Reward delivery evokes a burst of dopamine (DA) release in the nucleus accumbens (NAc), helping to promote reward-seeking behaviors^15,16^. The amplitude of this DA burst integrates multiple attributes of the reward, including both the reward’s size and how much effort went into it^17–19^. Although DA release is largely driven by the firing of DA neurons in the midbrain, studies have proposed a key role for local modulation of DA axon terminals in the striatum as well^20–23^. Which patterns of DA release are behaviorally relevant, what inputs determine these patterns, and whether these inputs dissociate DA cell body activity from striatal DA release, are all subject to debate^24–26^.

Recently, we found that DA release scales with preceding effort even if DA axon terminals in the NAc are stimulated directly via optogenetics^17^. When mice work for an identical optogenetic stimulation, more effort leads to more DA release. We reasoned that this variability in DA release might result from local modulation of DA axon terminals, a phenomenon that has been studied at length *in vitro*^20–23^ but has proven more difficult to isolate *in vivo*. Of the many modulators with potential to calibrate DA release, acetylcholine (ACh) consistently emerges as a potent effector, although the nature of this interaction has been contested^6–14,27^. In particular, there is ongoing debate over whether cholinergic interneurons in the striatum, which are capable of eliciting axonal DA release independent of DA cell body activity^6–11^, actually do so in any behavioral context^12–14^.

### Reward-evoked DA release encodes expended effort

To discover how reward-evoked DA release is modulated by effort, we employed a behavioral task^17^ that varies effort requirements for the same reward (Fig. 1a-b). Mice were first trained to nose poke during fixed-ratio 1 (FR1) and FR5 schedules of reinforcement for sucrose reward. Once they achieved accurate and stable responding, mice proceeded to a task in which effort was varied through descending 10-min FR blocks. Mice performed the task well, with very few inactive pokes (Extended Data Fig. 1a) and consistent behavior from day to day (Extended Data Fig. 1b and c). As the effort requirement increased, mice modulated their performance in the expected ways, initially escalating their poking rates to maintain a high level of reward consumption before reducing their poking rates and reward earnings (Extended Data Fig. 1d and e). Compared to low-effort rewards, rewards delivered after high effort were retrieved more quickly and consistently, suggesting high task engagement (Extended Data Fig. 1f and g).

**Fig 1.**
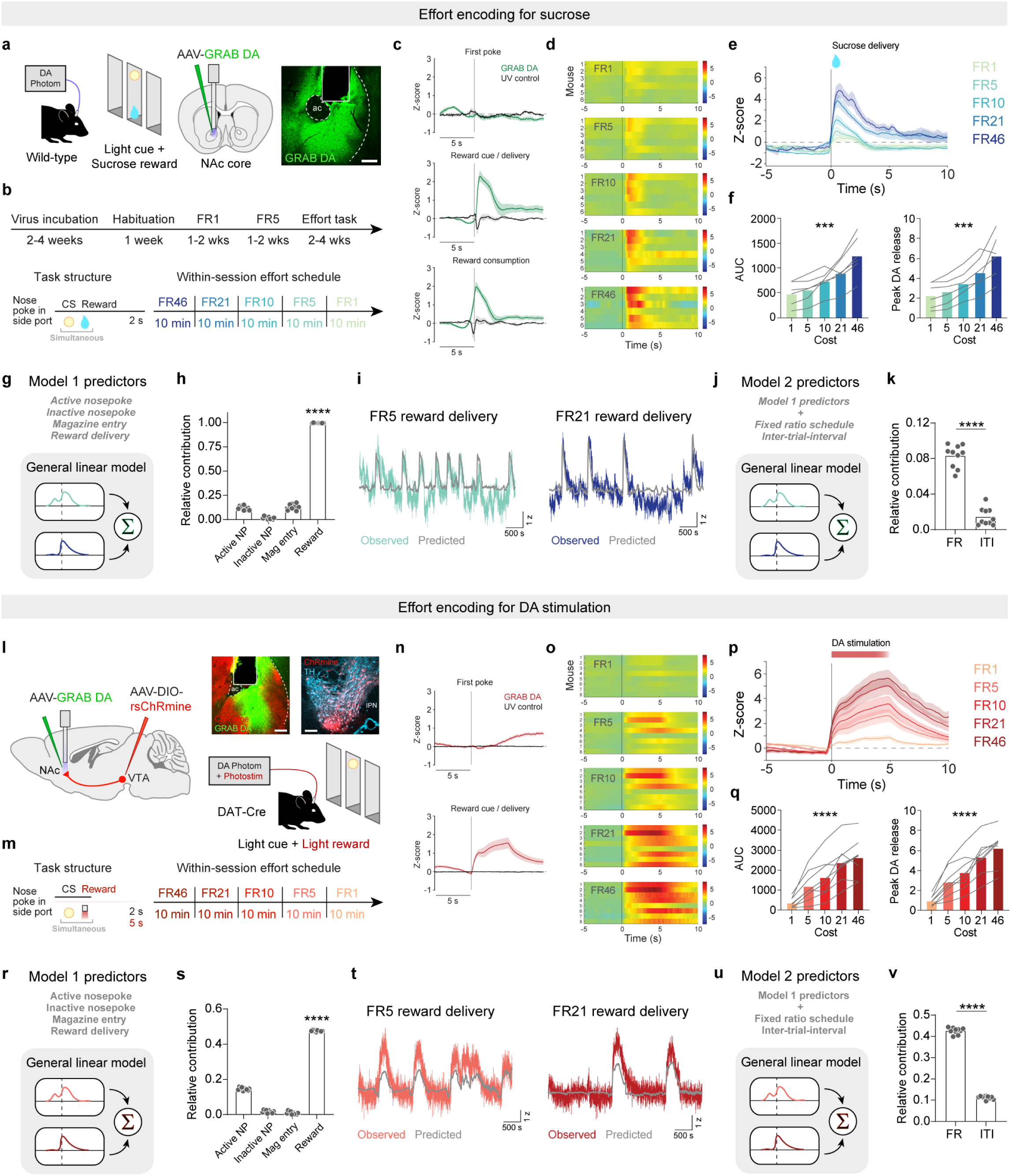
Reward-evoked DA release encodes expended effort. **a.** Schematic of effort task for sucrose reward and representative image of GRAB DA recording. Scale = 100 µm. **b.** Top, operant training schedule. Bottom left, task structure for obtaining rewards in the effort task. Bottom right, within-session schedule of effort blocks. Mice worked for rewards in 10 min blocks of FR46, FR21, FR10, FR5, and FR1. FR = fixed-ratio. **c.** DA release in NAc during the sucrose effort task aligned to the first nosepoke of each trial, reward delivery, and reward consumption, averaged across mice (*n* = 6). **d.** Heat maps illustrating the GRAB DA responses for each mouse across the FR schedules in the sucrose effort task. **e.** Average DA release aligned to the reward delivery for each FR block. **f.** Left, DA release area under the curve (0-4 s) for each FR. Friedman test, Friedman statistic = 23.33, ***p = 0.0001. Right, peak DA release for each FR. Friedman test, Friedman statistic = 23.33, ***p = 0.0001. **g.** Schematic of the general linear model used to predict DA release dynamics in the task. **h.** Contribution of each behavioral predictor to the total model R^2^, assessed using 10-fold cross-validation. One-way ANOVA, F(3,27) = 5744, ****p < 0.0001. Sidak-corrected multiple comparisons: active nosepoke vs inactive nosepoke, ****p < 0.0001; active nosepoke vs magazine entry, p = 0.12; active nosepoke vs reward delivery, ****p < 0.0001; inactive nosepoke vs magazine entry, ****p < 0.0001; inactive nosepoke vs reward delivery, p < 0.0001; reward delivery vs magazine entry, ****p < 0.0001. **i.** Actual versus model-predicted GRAB DA signal for example low- and high-effort trials. **j.** Schematic of model used to test the unique contributions of FR and ITI. **k.** Contribution of FR and ITI to the total model R^2^. Paired t-test, t = 11.39, ****p < 0.0001. **l-v**. Similar to **a-k**, except for optogenetic self-stimulation task. *n* = 8 DAT-Cre mice. Statistics in **q**: Left, Friedman test, Friedman statistic = 29.10, ****p < 0.0001. Right, Friedman test, Friedman statistic = 30.50, ****p < 0.0001. Statistics in **s**: One-way ANOVA, F(3,27) = 16250, ****p < 0.0001. Sidak-corrected multiple comparisons: active nosepoke vs inactive nosepoke, p < 0.0001; active nosepoke vs magazine entry, ****p < 0.0001; active nosepoke vs reward delivery, p < 0.0001; inactive nosepoke vs magazine entry, **p = 0.0025; inactive nosepoke vs reward delivery, ****p < 0.0001; reward delivery vs magazine entry, ****p < 0.0001. Statistics in **v**: Paired t-test, t = 67.81, ****p < 0.0001.

As mice worked for rewards, we recorded DA release in the NAc with GRAB DA (Fig. 1a). We observed robust DA release time-locked to reward delivery and consumption, but not during nosepokes (Fig. 1c). Consistent with our previous findings^17^, DA release at the time of reward delivery scaled with FR, such that high-effort sucrose rewards evoked more DA release (Fig. 1d-f). This increase in DA release at higher FRs was not due entirely to longer intervals between rewards, because we observed similar modulation regardless of inter-trial interval (ITI, Extended Data Fig. 2a-h).

To more precisely determine what aspects of the task were encoded by DA, we trained a general linear model (GLM) on various behaviors and task characteristics (kernels) (Fig. 1g). The GLM revealed a major contribution of reward delivery, but not nosepokes or magazine entries, to DA release (Fig. 1h). The model accurately predicted DA dynamics at both low and high efforts (Fig. 1i), with FR being a stronger predictor of DA release than ITI (Fig. 1j-k).

To determine if the effort encoding we observed for sucrose reward generalized to other types of reinforcers, we employed the same task but with optogenetic stimulation of DA axons in the NAc substituted for sucrose. DAT-Cre mice were prepared with Cre-dependent red-shifted ChRmine (rsChRmine) in the ventral tegmental area (VTA) and GRAB DA and an optical fiber in the NAc (Fig. 1l). These mice were subjected to the same task structure as for sucrose reward, except now for brain stimulation reward (Fig. 1m). Mice performed accurately (Extended Data Fig. 1h) and consistently (Extended Data Fig. 1i and j), with similar effort dependence as in the sucrose task (Extended Data Fig. 1k and l). When mice nosepoked for optical stimulation of DA release (5 s of 625-nm light at 20 Hz, 6 mW), DA release was again time-locked to reward delivery (Fig. 1n).

As with sucrose, we observed that DA release scaled with effort, such that the same optogenetic stimulation led to more DA at higher FRs (Fig. 1o-q). This effect was not due to the time elapsed between rewards (Extended Data Fig. 2i-p). In addition, changes in DA release were not due to opsin desensitization or “run down” of optically-evoked release over time, as 50 min of regular stimulations (independent of mouse behavior) showed similar DA release across the entire session (Extended Data Fig. 2q-u). Just as for the sucrose data, we trained a GLM on the photometry results (Fig. 1r) and identified the same patterns: the reward delivery kernel was the major contributor to model performance (Fig. 1s), the model accurately predicted release dynamics across FRs (Fig. 1t), and FR was a stronger predictor of DA release than ITI (Fig. 1u-v). Collectively, these data replicate and extend our prior work^17^ and support the hypothesis that DA release encodes expended effort.

### Effort encoding by NAc DA does not require DA cell body activity

We next sought to characterize the mechanism underlying the effect of effort on DA release. Modulation of DA release could involve alterations at the level of cell bodies in the VTA, axon terminals in the NAc, or a combination of the two^22^. Indeed, recent work has suggested that cell body and terminal activity can be dissociated in reward contexts^28^, although this theory remains contested^29^. Consistent with DA neuron recordings in monkeys^18^, we hypothesized that as mice exerted effort for reward, DA cell bodies would become more excitable, leading to synchronized activity and augmented DA release at the time of reward delivery. We tested this idea by expressing AAV-DIO-GCaMP8m in the VTA of DAT-Cre mice and implanting a fiber in the same location for recordings of DA cell bodies as mice worked for sucrose (Fig. 2a). We detected enhanced reward-evoked DA cell body activity at higher FR schedules, both during reward delivery and reward consumption (Fig. 2b and c). However, unlike NAc DA release, the enhancement of DA cell activity appeared to asymptote around FR21, with no significant increase at FR46 (Fig. 2b and c).

**Fig 2.**
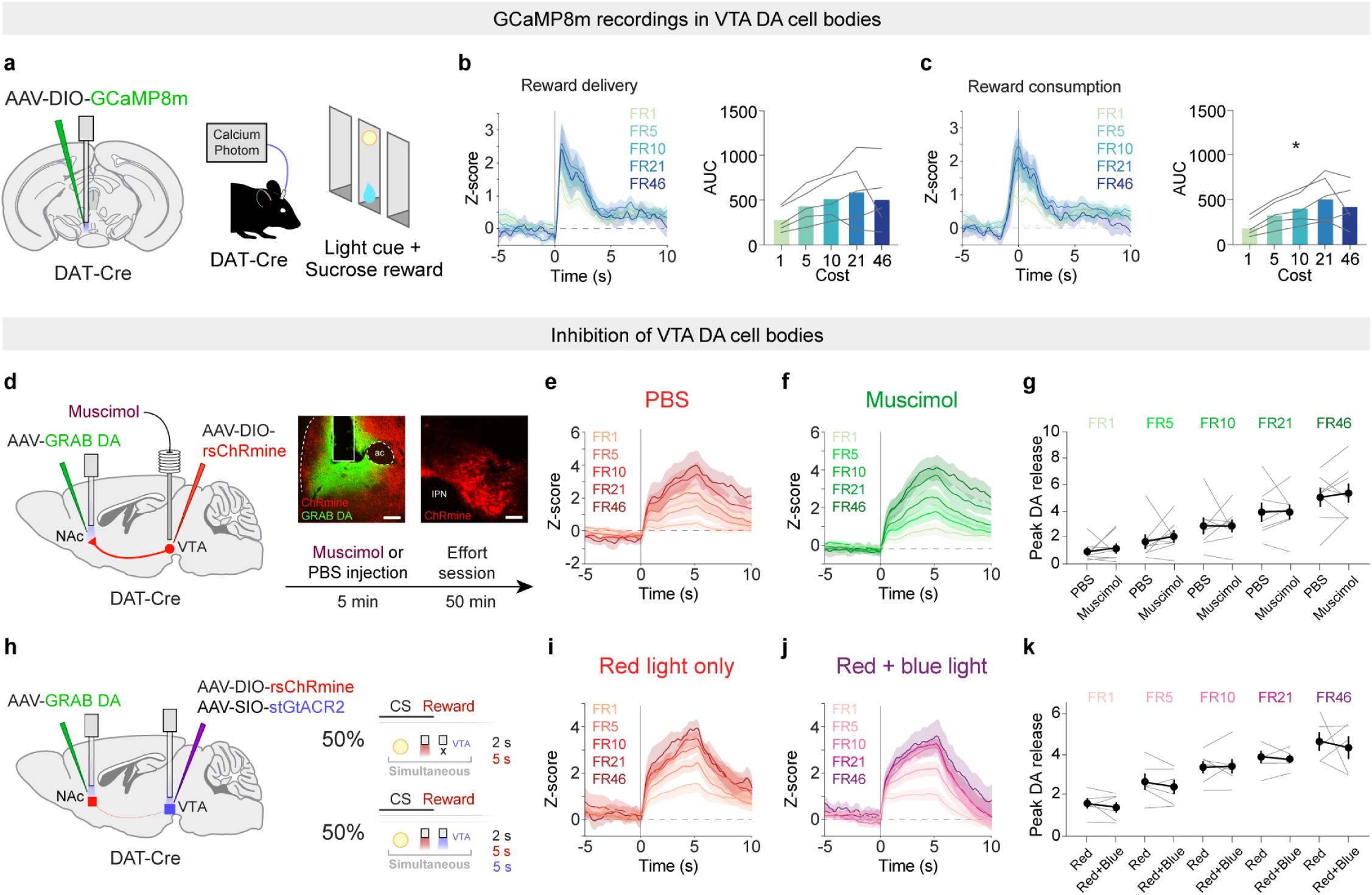
Effort encoding does not require VTA DA cell body activity. **a.** Schematic of VTA DA cell body recordings during effort task for sucrose reward. **b.** Left, average VTA GCaMP activity aligned to reward delivery (*n* = 5). Right, area under the curve (0-4 s) for each FR. Friedman test, Friedman statistic = 7.04, p = 0.13. **c.** Left, VTA GCaMP activity aligned to reward consumption. Right, area under the curve (0-4 s) for each FR. Friedman test, Friedman statistic = 12.16, *p = 0.016; *n* = 5. **d.** Left, schematic of AAV injection and cannula placement for inhibition of VTA cell bodies during the self-stimulation task. Top right, images of GRAB DA and rsChRmine expression in NAc and VTA. Bottom right, experimental schedule. Scale = 100 µm. **e.** DA release dynamics after microinjection of PBS into the VTA (*n* = 8). **f.** DA release dynamics after microinjection of muscimol into the VTA (*n* = 8). **g.** Peak DA (0-6 s) response for each FR for PBS and muscimol sessions. Mixed-effects model, fixed effect of FR, F(4,28) = 45.26, ****p < 0.0001; fixed effect of drug, F(1,7) = 0.24, p = 64; interaction of FR and drug; F(4,28) = 0.20; p = 0.93. Sidak-corrected multiple comparisons: FR1, PBS vs muscimol, p = 0.99; FR5, PBS vs muscimol, p = 0.97; FR10, PBS vs muscimol, p > 0.99; FR21, PBS vs muscimol, p > 0.99; FR46, PBS vs muscimol, p = 0.99; *n* = 8. **h.** Schematic of optogenetic inhibition of VTA DA cell bodies during self-stimulation task. **i.** DA release dynamics during control trials with no light in the VTA (*n* = 6). **j.** DA release dynamics during trials with blue light in the VTA to inhibit DA cell bodies (*n* = 6). **k.** Peak DA (0-6 s) response for each FR for trials with and without blue light in the VTA. Mixed-effects model, fixed effect of FR, F(4,37) = 26.10, ****p < 0.0001; fixed effect of light in the VTA, F(1,10) = 0.05, p = 0.82; interaction of FR and light in the VTA; F(4,37) = 0.66; p = 0.62. Sidak-corrected multiple comparisons: FR1, no light vs blue light, p > 0.99; FR5, no light vs blue light, p > 0.99; FR10, no light vs blue light, p > 0.99; FR21, no light vs blue light, p > 0.99; FR46, no light vs blue light, p = 0.77; *n* = 6.

These data suggest that modifications occurring in DA cell bodies may partially mediate the effort-based modulation of DA release in the NAc. To examine this possibility, we suppressed VTA DA cell body activity while recording NAc DA release as mice worked for rsChRmine stimulation of DA axon terminals. DAT-Cre mice were injected with GRAB DA in the NAc and DIO-rsChRmine in the VTA, and implanted with a fiber in the NAc and a unilateral guide cannula in the VTA for microinfusions of the GABA_A_ agonist muscimol to silence VTA neural activity (Fig. 2d). After training, mice were infused with muscimol or PBS into the VTA 5 min before an effort task session (Fig. 2d). Although unilateral muscimol successfully elicited the expected locomotor effects (Extended Data Fig. 3a-d), surprisingly the drug had no effect on DA effort-encoding (Fig. 2e-g).

Because muscimol inhibited the entire VTA, not only the DA cell bodies, we next took an optogenetic approach to selectively silence DA cell bodies. In DAT-Cre mice, we expressed GRAB DA in the NAc and infused a cocktail of Cre-dependent rsChRmine and the soma-targeted blue-light-sensitive anion channel GtACR2 (stGtACR2)^30^ in the VTA (Fig. 2h). We then implanted a fiber over the NAc for GRAB DA recordings and rsChRmine stimulation, and another fiber over the VTA for GtACR2-mediated inhibition of DA cell bodies. During the effort task, 50% of red-light rewards in the NAc were randomly paired with simultaneous blue-light exposure (5 s of 465 nm continuous light, 6 mW) in the VTA (Fig. 2h). Similar to muscimol, optical silencing of VTA DA cell body activity had minimal effect on effort-encoding in the NAc (Fig. 2i-k). This null effect could not be explained by nonfunctional opsins, because in the same mice, we found that red light alone triggered robust DA release (Extended Data Fig. 3e-h) while blue light alone potently inhibited DA release in the NAc (Extended Data Fig. 3i-k). Taken together, these data suggest that while effort encoding can be detected in VTA DA cell body activity, this activity is not necessary for the scaling of NAc DA release by effort.

### NAc nicotinic receptor signaling gates DA release associated with high effort

Since VTA DA cell body activity appears dispensable for effort-encoding in the NAc, we reasoned that physiological changes in DA axon terminals might support this phenomenon instead. As DA terminals in striatum are sensitive to various neuromodulators^22^, we turned to pharmacology to test the role of different modulators and GPCRs in DA effort-encoding. We prepared new DAT-Cre mice with rsChRmine in the VTA, GRAB DA in the NAc, and an optical fiber-coupled cannula implant in the NAc for drug microinfusions into the recording site (Fig. 3a and b and Extended Data Fig. 4a). We tested whether 9 different antagonists could disrupt effort-encoding of optogenetically-evoked DA release. Mice were trained in the task, habituated to microinjection procedures, and then tested for effort-encoding 5 min after microinfusions (Extended Data Fig. 4b). Intra-NAc blockade of alpha 1 adrenergic, muscarinic (pan and M5 specific), or neurokinin 3 receptors failed to affect DA effort-encoding, as did systemic blockade of A1 adenosine, A2A adenosine, neurokinin 1, or neurotensin 1 receptors (Extended Data Fig. 4c-l). In contrast, the nicotinic receptor antagonist dihydro-β-erythroidine (DHβE), which targets ⍺4β2 receptors, entirely blocked the enhanced DA release in high-FR blocks (Fig. 3c-f). Importantly, DHβE did not abolish light-evoked DA release altogether; rather, it prevented the increased release during high-effort rewards (Fig. 3e and f). Furthermore, DHβE was similarly effective at blunting DA effort encoding when mice worked for sucrose instead of optogenetic reward (Fig. 3g-l). Together, these data reveal ACh signaling at ⍺4β2 nicotinic receptors as a key and selective component of effort-encoding by DA.

**Fig 3.**
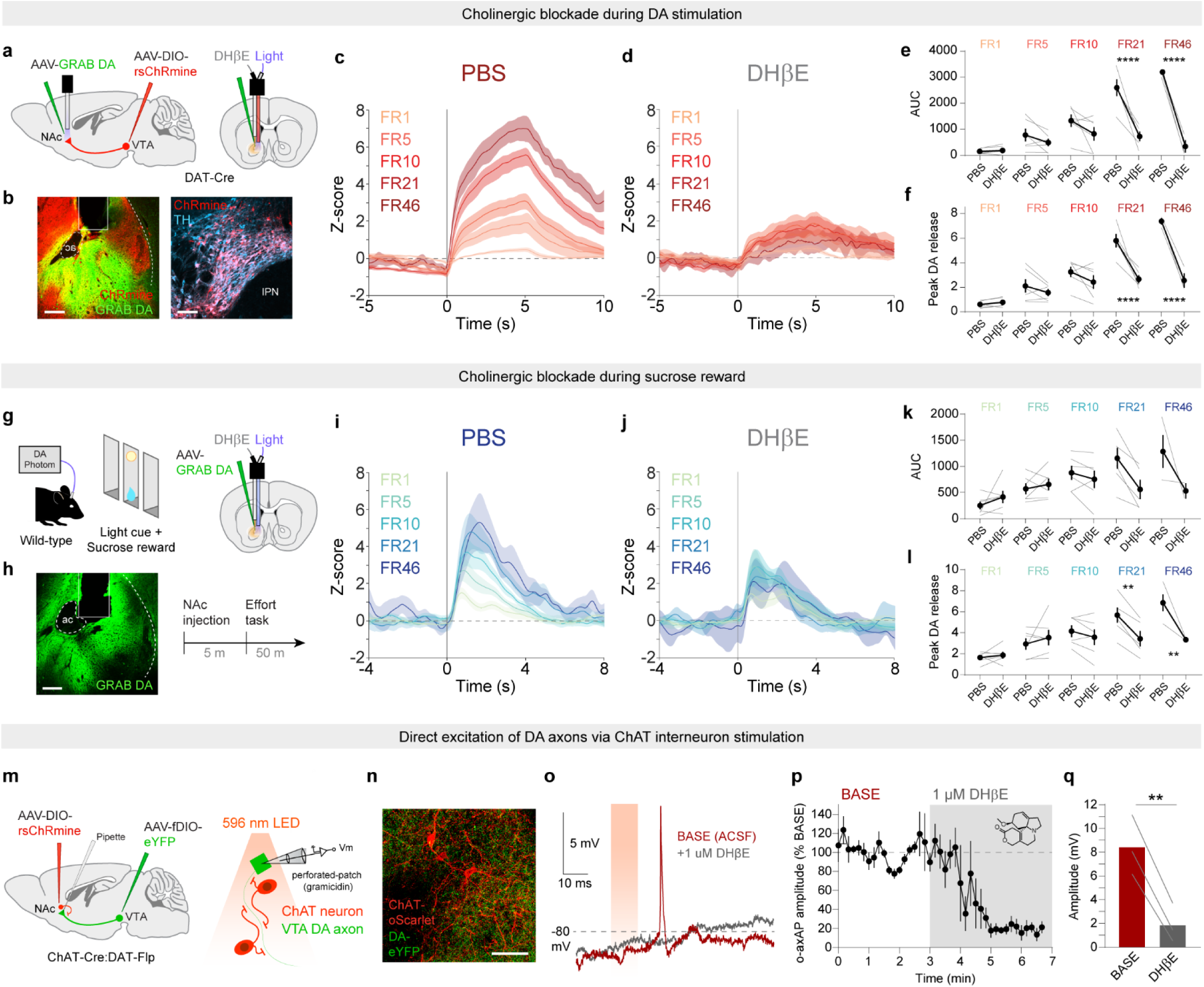
NAc nicotinic receptor signaling gates DA release during high effort. **a.** Joint optic fiber-cannula implants were used to locally infuse drugs prior to the self-stimulation task. **b.** Representative images of viral expression and fiber placement in the NAc and VTA. Scale = 100 µm. **c.** DA release across FR blocks during the self-stimulation task after microinjection of PBS into the NAc. *n* = 6. **d.** DA release across FR blocks during the self-stimulation task after microinjection of DHβE into the NAc. *n* = 6. **e.** Area under the curve (0-6 s) for each FR for PBS and DHβE sessions. Mixed-effects model, fixed effect of FR, F(4,20) = 20.63, ****p < 0.0001; fixed effect of drug, F(1,5) = 37.91, **p = 0.0016; interaction of FR and drug; F(4,16) = 24.43, ****p < 0.0001. Sidak-corrected multiple comparisons: FR1, PBS vs DHβE, p > 0.99; FR5, PBS vs DHβE, p = 0.78; FR10, PBS vs DHβE, p = 0.25; FR21, PBS vs DHβE, ****p < 0.0001; FR46, PBS vs DHβE, ****p < 0.0001, *n* = 6. **f.** Peak DA response (0-6 s) for each FR for PBS and DHβE sessions. Mixed-effects model, fixed effect of FR, F(4,20) = 39.33, ****p < 0.0001; fixed effect of drug, F(1,5) = 52.20, ***p = 0.0008; interaction of FR and drug; F(4,16) = 17.68; ****p < 0.0001. Sidak-corrected multiple comparisons: FR1, PBS vs DHβE, p > 0.99; FR5, PBS vs DHβE, p = 0.77; FR10, PBS vs DHβE, p = 0.36; FR21, PBS vs DHβE, ****p < 0.0001; FR46, PBS vs DHβE, ****p < 0.0001; *n* = 6. **g-l.** Similar to **a-f**, except for the sucrose task. Statistics for **k**: Mixed-effects model, fixed effect of FR, F(4,20) = 6.54, **p = 0.0016; fixed effect of drug, F(1,5) = 2.85, p = 0.15; interaction of FR and drug; F(4,14) = 3.83; *p = 0.027. Sidak-corrected multiple comparisons: FR1, PBS vs DHβE, p = 0.94; FR5, PBS vs DHβE, p > 0.99; FR10, PBS vs DHβE, p = 0.98; FR21, PBS vs DHβE, p = 0.062; FR46, PBS vs DHβE, p = 0.11; *n* = 6. Statistics for **l**: Mixed-effects model, fixed effect of FR, F(4,20) = 11.75, ****p < 0.0001; fixed effect of drug, F(1,5) = 14.71, *p = 0.012; interaction of FR and drug; F(4,14) = 6.56, **p = 0.0034. Sidak-corrected multiple comparisons: FR1, PBS vs DHβE, p > 0.99; FR5, PBS vs DHβE, p = 0.75; FR10, PBS vs DHβE, p = 0.80; FR21, PBS vs DHβE, **p = 0.0058; FR46, PBS vs DHβE, **p = 0.0035; *n* = 6. **m.** Schematic of AAV injection and electrophysiological recording configuration. **n.** Representative brain slice image. Scale bar = 50 µm. **o.** Representative axonal AP traces from an eYFP+ VTA DA axon bleb evoked by rsChRmine stimulation of cholinergic interneurons at baseline and in the presence of 1 µM DHβE. **p.** Time course of average axonal APs evoked by rsChRmine stimulation before and during acute perfusion of 1 µM DHβE; *n* = 3 axons, N = 3 mice **q.** Average o-axAP amplitude over 10 sweeps in baseline and 1 µM DHβE conditions, Paired t-test, t = 12.54, df = 2, **p = 0.0063, *n* = 3 axons, N = 3 mice.

Interactions between DA and ACh are well-established^31–35^, with substantial evidence *in vitro* that ACh can directly excite DA axons^7,11^ and trigger DA release^6–11^ through nicotinic receptors in the dorsal striatum. To determine if these dorsal striatal findings generalize to the NAc, we crossed DAT-Flp mice to ChAT-Cre mice and injected Flp-dependent eYFP into the VTA and Cre-dependent rsChRmine into the NAc. Acute slices were prepared and perforated patch recordings of DA axons within the NAc were performed while stimulating cholinergic interneurons with red light (Fig. 3m and n). We detected robust excitation of DA axons in response to optogenetic stimulation of local cholinergic interneurons, including excitatory postsynaptic potentials (EPSPs), single action potentials (APs), compound AP-EPSPs, and double APs (Fig. 3o and Extended Data Fig. 5) that were blocked by 1 µM DHβE (Fig. 3p-q and Extended Data Fig. 5). These data confirm that cholinergic interneurons can drive DA release locally in the NAc.

### NAc ACh release encodes expended effort

If ACh gates DA release to encode effort, then ACh release dynamics should also reflect exerted effort. We therefore prepared new wild-type mice with GRAB ACh in the NAc and ran them through the same behavioral procedures (Fig. 4a and b). Consistent with previous work^7,36–38^, we observed complex ACh release waveforms during different phases of the task (Fig. 4c). Focusing on the response around reward delivery, we observed a subtle ramping of activity directly prior to reward, followed by a triphasic ACh waveform with a sharp peak right after delivery, a rapid dip, and a second peak around the time of reward consumption (Fig. 4c). These dynamics were robustly modulated by FR (Fig. 4d-f), even after controlling for the effect of ITI (Extended Data Fig. 6a-i). Bleaching of the photometry signal over the course of the session cannot explain our observations, since we found the same effort encoding in a control session with ascending FR blocks (Extended Data Fig. 6j-m). Finally, ACh appeared to track effort but not reward size, because in a separate task that kept effort constant but varied sucrose concentration between 5% and 32%, ACh release was nearly identical for both rewards (Extended Data Fig. 6n-w). These data suggest that, contrary to DA release, which tracks both reward size and effort^17^, ACh release in the NAc tracks effort alone.

**Fig 4.**
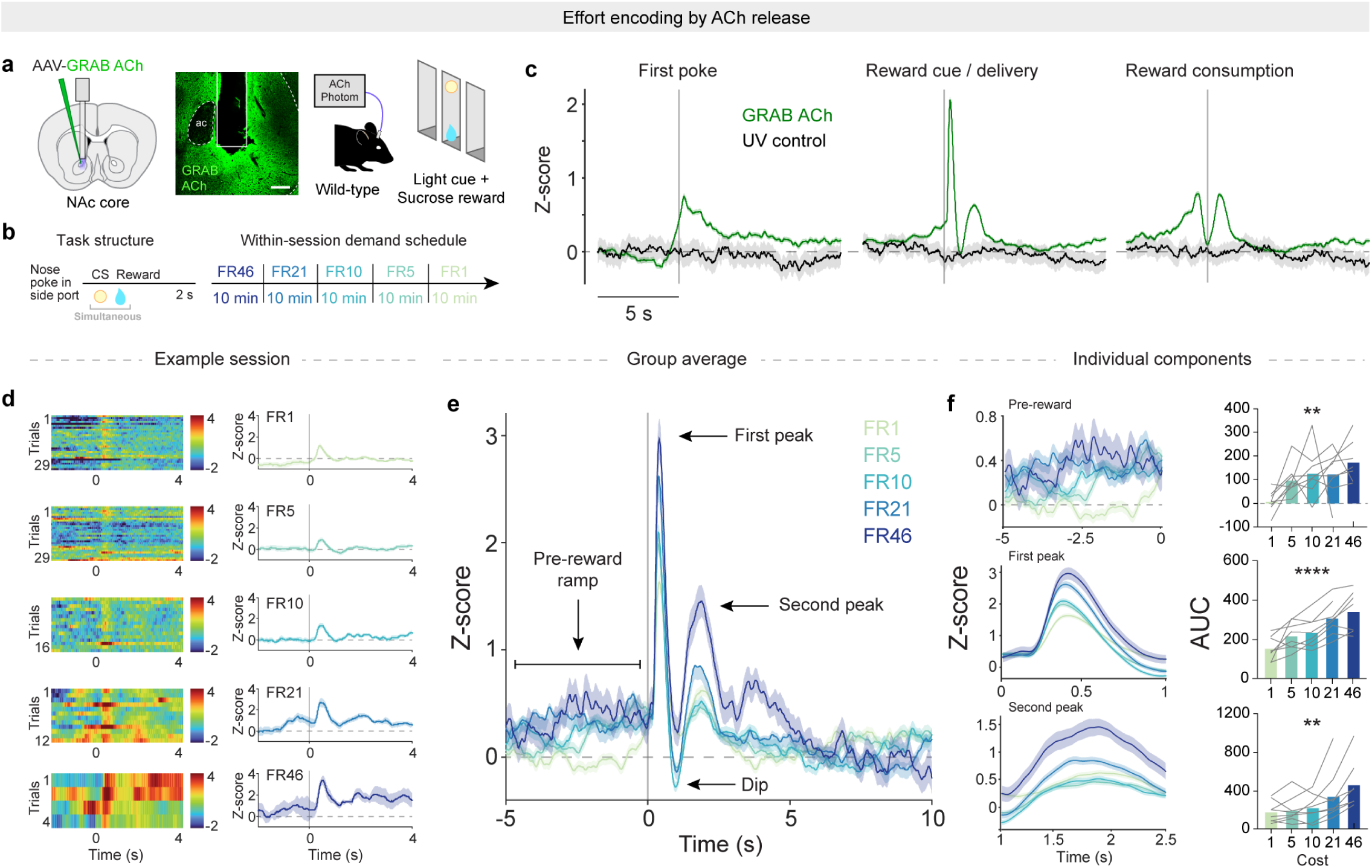
ACh release encodes expended effort. **a.** Schematic of ACh recordings during sucrose task, with representative image of GRAB ACh expression. Scale = 100 µm. **b.** Left, trial structure. Right, within-session schedule of effort blocks. **c.** ACh release (green) and UV control signal (black) in NAc during the sucrose task aligned to the first nosepoke, reward delivery, and reward consumption. *n* = 8. **d.** Heat maps and traces of reward-evoked ACh release from an example session. **e.** ACh release across FRs time-locked to reward delivery, averaged across mice (*n* = 8). **f.** Expanded view of individual components of ACh release waveform. Top left, pre-reward ramp. Top right, area under the curve of the pre-reward period (-5-0 s) for each FR. Mixed-effect model, fixed effect of FR, F(4,34) = 4.47, **p = 0.0052. Center left, first peak. Center right, area under the curve of the first peak (0.25-0.75 s). Mixed-effect model, fixed effect of FR, F(4,27) = 19.64, ****p < 0.0001. Bottom left, second peak. Bottom right, area under the curve for the second peak (1-2.5 s). Mixed-effect model, fixed effect of FR, F(4,27) = 6.17, **p = 0.0012, *n* = 8.

While the NAc receives ACh inputs from multiple sources, local interneurons are thought to predominate when it comes to modulation of DA^39^. We recorded the activity of cholinergic interneurons in the NAc during the effort task in ChAT-Cre mice expressing Cre-dependent GCaMP8m (Extended Data Fig. 7a and b). We observed a strikingly similar set of dynamics to our GRAB ACh recordings (Extended Data Fig. 7c-f), again with effort encoding independent from ITI (Extended Data Fig. 7g-o). These data indicate that cholinergic interneurons in the NAc are activated during states of high effort, putting them in a position to potentiate DA release upon reward delivery.

### Local cholinergic signaling is essential for DA effort encoding and effortful behavior

To examine how ACh might augment DA release in this task, we first inspected the timing of ACh and DA release relative to reward delivery (Extended Data Fig. 8). Direct comparison suggested that reward-evoked ACh release precedes DA release by roughly 400 msec, a lag that holds true across all FR blocks, even though the on-kinetics of the GRAB DA sensor are faster than those of GRAB ACh (personal communication, Y. Li).

This temporal offset supports our hypothesis that an initial burst of ACh gates DA release to high-effort rewards, but the evidence is correlational. To directly test the role of ACh in DA effort-encoding, we prepared new ChAT-Cre mice with GRAB DA and Cre-dependent NpHR, an inhibitory opsin sensitive to red light, in the NAc (Fig. 5a). These mice were trained to perform the sucrose task and then subjected to alternating days of red light exposure for 4 s after each reward delivery (Fig. 5b). In trials with no red light, we observed the characteristic effort-encoding by DA. When red light was delivered, however, effort-encoding was significantly blunted (Fig. 5c and d). Similar to blockade of nicotinic receptors, we found a progressive disruption of DA release as effort increased, where low-effort DA release was no different than control conditions (Fig. 5e and f). We found no such disruption of DA effort-encoding in control mice expressing mCherry rather than NpHR in cholinergic neurons (Extended Data Fig. 9a-f).

**Fig 5.**
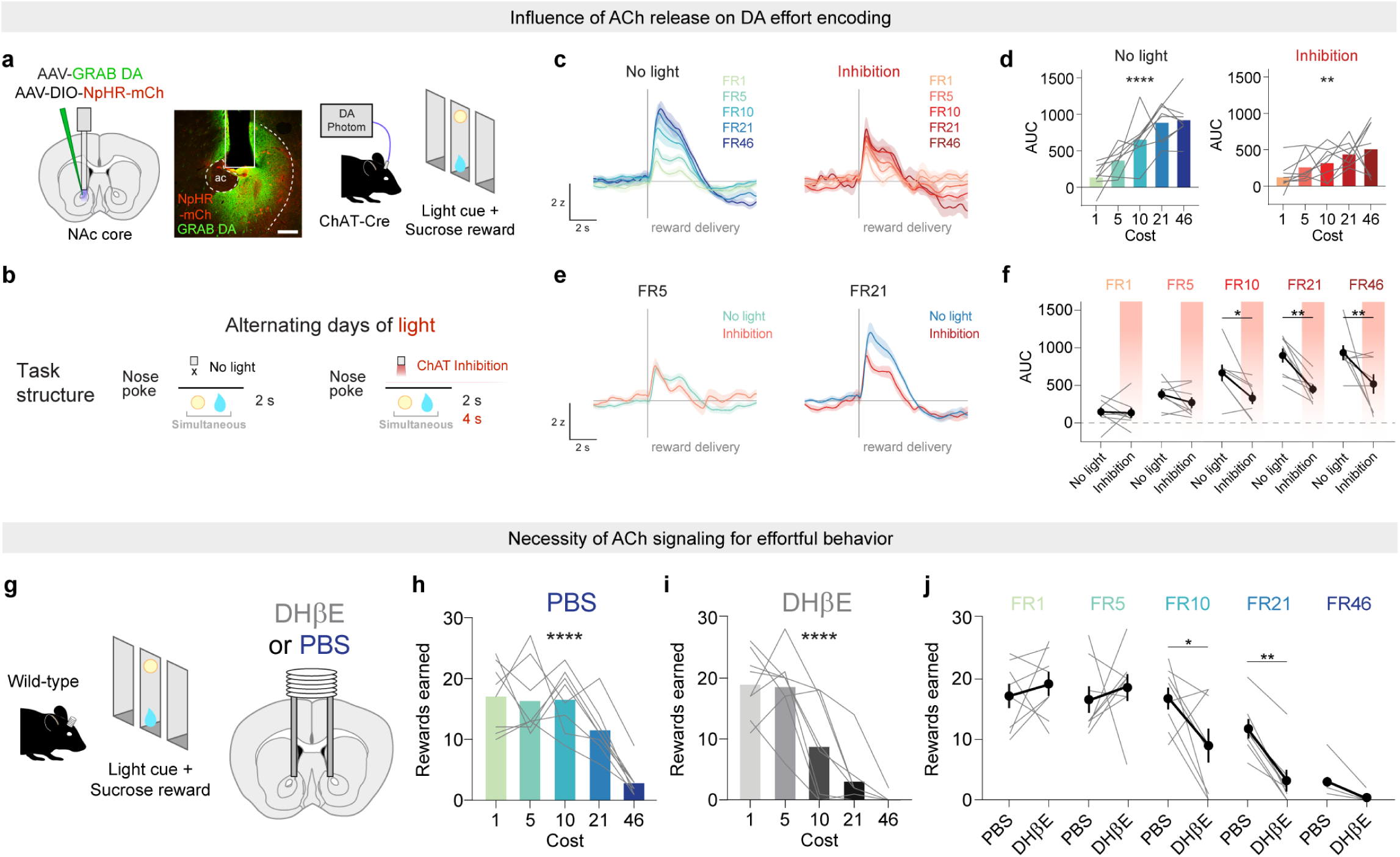
Local cholinergic signaling is essential for DA effort encoding and effortful behavior. **a.** Schematic and representative image of DA recordings with cholinergic neuron inhibition during the sucrose task. Scale = 100 µm. **b.** Experimental design and task structure with alternating days of red light to inhibit cholinergic neurons. **c.** DA release dynamics during the sucrose task for control sessions with no red light (left) and sessions with red light inhibition of cholinergic neurons (right). *n* = 8 mice. **d.** Left, quantification of area under the curve (0-4 s) for each FR during no light sessions. Friedman test, Friedman statistic = 25.10, ****p < 0.0001. Right, area under the curve during red light sessions. Friedman test, Friedman statistic = 16.40, **p = 0.0025, *n* = 8. **e.** Direct comparison of reward-evoked DA release in sessions with red light off or on for FR5 trials (left) and FR21 trials (right). *n* = 8. **f.** Area under the curve (0-4 s) for each FR directly comparing the effect of optical inhibition. Mixed-effects model, fixed effect of FR, F(4,28) = 21.94, ****p < 0.0001; fixed effect of inhibition, F(1,7) = 23.05, **p = 0.0020; interaction of FR and inhibition; F(4,28) = 3.52, *p = 0.019. Sidak-corrected multiple comparisons: FR1, no light vs inhibition, p = 0.99; FR5, no light vs inhibition, p = 0.96; FR10, no light vs inhibition, *p = 0.024; FR21, no light vs inhibition, **p = 0.0012; FR46, no light vs inhibition, **p = 0.0043, *n* = 8. **g.** Schematic of sucrose task and bilateral cannula implant in NAc for PBS or DHβE infusion. **h.** Rewards earned at each FR after microinjection of PBS. Friedman test, Friedman statistic = 21.25, ****p < 0.0001, *n* = 8. **i.** Rewards earned at each FR after microinjection of DHβE. Friedman test, Friedman statistic = 25.33, ****p < 0.0001, *n* = 8. **j.** Rewards earned for each FR directly comparing performance after PBS or DHβE infusion. Mixed-effects model, fixed effect of FR, F(4,28) = 36.36, ****p < 0.0001; fixed effect of drug, F(1,7) = 5.26, p = 0.056; interaction of FR and drug; F(4,28) = 4.85, **p = 0.0042. Sidak-corrected multiple comparisons: FR1, PBS vs DHβE, p = 0.95; FR5, PBS vs DHβE, p = 0.90; FR10, PBS vs DHβE, *p = 0.016; FR21, PBS vs DHβE, **p = 0.0069; FR46, PBS vs DHβE, p = 0.79, *n* = 8.

One possible consequence of increased DA release to high-effort rewards is the maintenance of effortful responding. Thus, we tested if ACh-DA interactions in the NAc promote effortful behavior. We prepared new mice with bilateral guide cannulas targeting the NAc for bilateral DHβE infusions (Fig. 5g). Compared with PBS infusions, DHβE treatment reduced sucrose reward responding and increased latency to initiate trials at high FR blocks but not low FR blocks (Fig. 5h-j and Extended Data Fig. 9g). To control for drug metabolism across the session, we tested the same mice in a counterbalanced, across-session schedule where mice worked for sucrose at either FR1 or FR21 on alternating days (Extended Data Fig. 9h and i). Infusion of DHβE had no effect at FR1 but strongly reduced reward consumption at FR21 (Extended Data Fig. 9j and k). These findings support the proposition that when rewards are earned through high effort, ACh signaling potentiates DA release in the NAc, promoting continued effort exertion.

## Discussion

We found that in high-effort conditions, ACh levels rise quickly in response to reward delivery, binding to α4β2 nicotinic receptors on DA axons to augment DA release. This increase in DA release then motivates continued effort expenditure, because blocking the nicotinic receptors on DA axons reduced reward earnings specifically when high effort was required.

Our results help explain an observation across the animal kingdom that individuals value rewards more when they have worked harder for them^2–5,40–43^. Such a process may be adaptive in promoting reward-seeking even when costs are high or rewards are scarce. We found that this evolutionarily-conserved psychological effect appears to depend on the same receptors responsible for the reinforcing effects of nicotine^44^. Future mechanistic models of addiction, therefore, may benefit from considering how the added value of effort, mediated by local interactions between ACh and DA axons, may reinforce drug-seeking behavior.

*In vitro* studies have long demonstrated robust cholinergic regulation of DA release^6–11^. In slice, the stimulation of even a single cholinergic interneuron can elicit DA release in a wide swath of the striatum^6^. *In vivo* studies, however, have mostly failed to find large effects of ACh on DA release, at least in baseline conditions or during simple motor or reward tasks^12–14^. But these *in vivo* studies did not examine ACh-DA interactions in high-effort contexts. Only by modulating effort requirements were we able to unmask a functionally-relevant effect of ACh on DA release. Our findings imply that that the interactions between neuromodulators are highly contingent on behavioral context.

The findings are also consistent with studies showing decreased motivation^27^ and DA release^7,45^ with blockade or deletion of nicotinic receptors. It is important to note, however, that the role of ACh in modulating DA and motivation is complex, with influential work demonstrating that cholinergic interneurons can oppose, rather than enhance, the motivational influence of appetitive cues^46–50^. Our results here are temporally precise, at the time of reward delivery, and limited to the contribution of nicotinic receptors on DA axons. It is likely that ACh-DA interactions vary depending on the ongoing behavior^51^, the preceding activity of DA and ACh neurons^22,23,50^, the region of striatum^37,50^, and the type of ACh receptor examined^49^.

Besides shedding light on effort valuation and the interactions between DA and ACh, our work speaks to an ongoing debate about whether the activity of DA cell bodies can be dissociated from the release of DA in striatal targets^28,29^. We found that in the case of sucrose rewards, VTA DA neurons encode expended effort (Fig 2a-c), likely accounting for at least some of the effect we observed on DA release in the NAc. At the same time, however, inhibiting VTA DA neurons did not reduce effort encoding in the NAc (Fig. 2d-k), while blocking nicotinic receptors on DA axon terminals did (Fig 3a-l). Thus we show that in the right context, local ACh can expand the dynamic range of DA release beyond what is driven by cell body activity, with an important role in boosting effortful behavior.

## Materials and Methods

### Subjects

Male and female mice of C57BL/6J background (Jackson Laboratories, strain #000664) were used in approximately equal numbers for all experiments. Wild-type mice were crossed with DAT:IRES-Cre mice (Jackson Laboratory, strain #006660) or ChAT:IRES-Cre mice (Jackson Laboratory, strain #006410). For slice electrophysiology experiments, ChAT:IRES-Cre mice were crossed with DAT-Flp mice (Jackson Laboratories, strain #035436). All transgenic animals used in experiments were genotyped and found to be heterozygous for Cre- or Flp-recombinase. Mice were separated by sex and group-housed after weaning before surgical procedures or behavioral assays. All behavioral experiments were conducted during the dark cycle (12 hour light/dark) when mice were 10-24 weeks old. Mice were housed at ∼21C in 30-70% humidity, and they were food restricted to 85% of *ad libitum* body weight for all behavioral experiments to facilitate motivated behavior. All procedures complied with the animal care standards set forth by the National Institutes of Health (NIH) and were approved by the Stanford University Administrative Panel on Laboratory Animal Care.

### Behavioral training

Food restriction and behavioral training began at a minimum of 2 weeks after virus injection and fiber implantation. Prior to starting behavioral experiments, mice were habituated for 2 days, including handling, tethering to the patch cords, and allowing exploration of the operant boxes (Med Associates) for 30 minutes. After habituation, mice were exposed to 1 day of a Pavlovian task in which rewards were delivered at random intervals spanning 25-35 s for 30 minutes.

Operant chambers were equipped with a speaker for white-noise and three identical nosepoke ports, each with an associated cue-light and an infrared emitter-sensor to measure port entry times. Rewards were delivered in conjunction with a 2 s cue comprising white noise and central port light so that mice would associate reward delivery with the light-sound cue. For sucrose rewards, 10µL of 32% (w/v) sucrose was used, apart from one experiment in which 5% sucrose was used (Extended Data Fig. 6n-w). After 1 day of Pavlovian training, mice progressed to the operant task. Mice trained on FR1 for a minimum of 5 sessions. The active nosepoke port (left or right) was counterbalanced between mice; each mouse continued with the same active port for all experiments. On days 1-3, the active food port was baited with a crushed portion of fruit loop to encourage exploration of that port. After earning at least 10 rewards at FR1, with an accuracy (% active nosepokes) >80%, mice progressed to FR5. Mice continued FR5 for at least 5 sessions and until the number of rewards earned per session remained within 20% for 3 consecutive days. Once this criterion was met, mice progressed to the sucrose effort task. We did not attempt to equalize reward consumption between mice, but rather aimed to ensure that all mice had accurate and internally consistent performance. Training for optogenetic self-stimulation was achieved in the same manner as for sucrose except the sucrose reward was replaced with 5 s optogenetic stimulation paired with the 2 s light and white noise cue. All behavioral tasks were coded in Med-PC V (Med Associates).

### Sucrose effort task

The sucrose effort task comprised five 10-minute blocks of fixed ratios including FR46, FR21, FR10, FR5 and FR1. After mice poked in the active port for the required number of times, sucrose reward (10µL 32% (w/v)) was delivered in a central magazine, accompanied by a 2 s light and white noise cue. Once the cue stopped and mice entered the magazine to consume reward, they were free to start the next trial at their own pace by poking in the active port. As in our prior work^17^, the fixed ratios were presented in descending fashion to prevent mice from achieving early satiety under low cost conditions, except for one control experiment with ascending presentation of blocks (Extended Data Fig. 6j-m). Block transitions were not signaled to the mice.

### Modified sucrose effort task

For microinjection experiments with PBS or DHβE, due to concern that the behavioral effect of the drug was a result of the drug wearing off over the course of the session, we employed a modified sucrose effort task. In the modified task, after PBS or DHβE injection, mice underwent a 30-min session in which the poking requirement was kept constant for the entire session at either FR5 or FR21. The order of the 4 experimental days (PBS vs DHβE; FR5 vs FR21) was counterbalanced between mice.

### ChRmine effort task

The ChRmine effort task was structured identically to the sucrose effort task, except that mice worked for optogenetic DA stimulation. ChRmine stimulation was paired with the same 2 s light and white noise cue in the magazine port, and upon cue cessation, the mice were free to start the next trial at their own pace.

### Stereotactic injections and viruses

Mice (8-12 weeks old) were anesthetized with isoflurane (5% induction, 1-2% maintenance). Subjects were fixed on a stereotaxic frame (Kopf Instruments), a small incision was made, and burr holes were drilled in the skull over the sites of injection or fiber implantation. The following coordinates relative to bregma were used: VTA, -3.1 mm anteroposterior (AP), ±0.4 mm mediolateral (ML), 4.2 mm dorsoventral (DV); NAc core, 1.5 mm AP, ±0.9 mm ML, 4.1 mm DV from the skull surface. Microliter syringes (Hamilton) were lowered to the specified depth from the skull and used to inject 500 nL of virus solution at a flow rate of 0.1-0.25 mL/min. Borosilicate optic fibers for photometry and/or optogenetic stimulation with 200-400 µm diameter and numerical aperture 0.66 (Doric) were implanted directly above the striatal or midbrain virus injection site and secured to the skull using screws (Antrin Miniature Specialties) and light-cured dental adhesive cement (DenMat, Geristore A&B paste). For the cohort with recordings in NAc and cannula or optogenetic fiber placement in the midbrain (Figure 2, S3), the VTA implant was cemented first and then the NAc fiber was implanted in the same hemisphere. Cannulas for drug microinfusion were implanted 1.5 mm above the target site with the injector extending to the site. For all cohorts, the hemisphere targeted for recordings was counterbalanced between mice. Adeno-associated viruses (AAVs) used for stereotactic injections were AAV9-hSyn-GRAB_DA_ DA2m (DA4.4, WZ Biosciences), AAV2/9-hSyn-gACh4m (GRAB ACh, Brain VTA), AAV5-hSyn1-SIO-stGtACR2-FusionRed (Addgene), AAV9-syn-FLEX-jGCaMP8m-WPRE (Addgene), AAV-8-EF1a-DIO-rsChRmine-oScarlet-WPRE (Stanford Gene Vector and Virus Core), AAV-dj-Ef1a-DIO-NpHR3.0-mCherry (Stanford Gene Vector and Virus Core), AAV-dj-Ef1a-DIO-eYFP (Stanford Gene Vector and Virus Core), AAV-dj-Ef1a-fDIO-eYFP (Stanford Gene Vector and Virus Core), and AAV-dj-Ef1a-DIO-mCherry (Stanford Gene Vector and Virus Core). AAV titers ranged from 1 x 10^12^ to 2 x 10^13^ gc/mL.

### Fiber photometry

Mice implanted with optical implants (400 µm diameter, 0.66 numerical aperture, Doric lenses) were connected via a ceramic sleeve (Precision Fiber Products) to low-autofluorescence patch cords of matching diameter and numerical aperture (Doric). Signals passed through a fiber optic rotary joint (Doric) before filtering through a fluorescence mini cube (Doric) and reaching a femtowatt photodetector (Newport, #2151). LEDs (Doric) emitted frequency-modulated ultraviolet (405 nm) and blue (465 nm) light to stimulate control and either GRAB or GCaMP signals through the same fiber. Blue LED power was adjusted to ∼35µW at the fiber tip and lowered as needed if signal saturated the photodetector. Ultraviolet LED power was adjusted to ∼10 µW at the fiber tip. Digital signals were sampled at 1.0173 kHz, demodulated, lock-in amplified, and recorded by a real-time signal processor (RZ5P, Tucker-Davis Technologies) using Synapse software (Tucker-Davis Technologies). Synchronized behavioral events from the Med Associates boxes were collected directly into the RZ5P digital input ports. To reduce any confounds from photobleaching, animals were recorded for about 5 min before behavioral testing began.

### Optogenetic manipulations

Optogenetic manipulations with rsChRmine and NpHR were conducted simultaneously as fiber photometry recordings through the same optical implants. Optogenetic manipulation with stGtACR was conducted through optical implants of 200 µm with 0.66 numerical aperture without simultaneous fiber photometry recording. Optogenetic manipulation with rsChRmine or NpHR was conducted by connecting a 625 nm LED light source (Prizmatix) via a plastic fiber (1mm diameter, NA 0.63) and a fiber optic rotary joint (Doric). rsChRmine stimulation was performed at a stimulation of 5 s, 20 Hz, and 6 mW with a pulse width of 10 ms. Optogenetic inhibition with NpHR or control mCherry was conducted with constant 4 s, ∼6 mW light. Optogenetic manipulation with stGtACR2 was conducted with a 450 nm LED light source (Prizmatix), using 5-10 s of constant ∼10 mW light.

### Drug administration

For systemic injection experiments, mice were administered intraperitoneal injections of aprepitant 10 mg/kg (Axon Medchem), DPCPX 2 mg/kg (Tocris), istradeffyline 2 mg/kg (Tocris), or SR 48692 5 mg/kg (Tocris). DPCPX and instradeffyline were dissolved in DMSO (Sigma-Aldrich) and diluted to 20% DMSO in 0.9% saline to prevent precipitation. Aprepitant and SR 48692 were dissolved in DMSO and diluted to 5% DMSO in corn oil (Sigma-Aldrich) to prevent precipitation. Mice were injected with a volume of 10 mL/kg. After injection, mice were placed in their home cages for 10 minutes, then placed in the operant chambers for 5 minutes of habituation prior to beginning the rsChRmine effort task. For these agents, systemic injection was chosen over intracranial injection due to risk of brain injury from the necessary solvents.

For local injection experiments, mice were administered an intra-VTA infusion of muscimol 1 mM (Tocris) or intra-NAc infusion of dihydro-β-erythroidine (DHβE) hydrobromide 30 µg/hemisphere (Tocris), terazosin 3 µg (Tocris), osanetant 375 ng (Axon Medchem), scopolamine 22 nmol (Tocris), or ML375 105 pmol (Axon Medchem). Osanetant was dissolved in Tween 20% until homogeneous, then diluted to 1% Tween in filtered molecular-quality PBS to prevent precipitation. All other antagonists were dissolved in filtered molecular-quality PBS only. Drugs were infused through an injector cannula connected to a 5 µL Hamilton syringe using a microinfusion pump (Harvard Apparatus) at a continuous rate of 150 nL per minute to a total volume of 0.3 µL unilaterally or bilaterally. For unilateral infusions with fiber photometry recording, multiple fluid injection cannulas (Doric) were used. For bilateral infusions, bilateral infusion cannulas (P1 Technologies) were used. Injector cannulas were removed 2 min after infusions were complete. After infusion, mice were placed in the operant chambers and allowed to habituate for 5 minutes prior to beginning the sucrose or rsChRmine effort task.

### Slice electrophysiology

Acute brain slices were prepared from adult ChAT-Cre:DAT-Flp mice between postnatal days P120-P180. ChAT-Cre:DAT-Flp mice were sterotaxically injected with 500 nL of AAVdj-eF1a-fDIO-eYFP into the VTA (AP: -3.1, ML: 0.4, DV: -4.2) and AAV8-Ef1a-DIO-rsChRmine-oScarlet into the NAc (AP: +1.5, ML: 0.9, DV: -4.1) at P90-P120 and were used for electrophysiology experiments 4-8 weeks post injection. Mice were anesthetized with isoflurane anesthesia and transcardially perfused prior to decapitation. Brains were carefully extracted and 250 µm thick coronal slices were cut using a vibratome (Leica VT1200 S) in ice-cold slice solution containing: 110 mM sucrose, 62.5 mM NaCl, 2.5 mM KCl, 6 mM MgCl_2_, 1.25 mM KH_2_PO_4_, 26 mM NaHCO_3_, 0.5 mM CaCl_2_, 20 mM D-glucose, pH 7.35-7.40. Slices were incubated in slice solution for 20 minutes at 33°C, then transferred to a room-temperature holding chamber with artificial cerebrospinal fluid (ACSF, 125 mM NaCl, 2.5 mM KCl, 1 mM MgCl_2_, 1.25 mM KH_2_PO_4_, 26 mM NaHCO_3_, 2 mM CaCl_2_, and 20 mM D-glucose, pH 7.35-7.40). Brain slices were stored in room temperature ACSF and used for recording 30 to 300 minutes later. All solutions were saturated with carbogen.

Individual brain slices were placed in an RC-27 recording chamber (Warner Instruments) and superfused with carbogen-saturated ACSF (33–36°C) at a flowrate of 2-3 mL/min. eYFP-positive cut ends of axons (blebs) originating from the VTA present at the slice surface in the NAc were visualized with a BX51WI upright microscope (Olympus) equipped with epifluorescence, IR-DIC optics, 470 nm (ET-GFP #49002, Chroma) and 596/83 nm single-band bandpass (FF01-596/83-25, Semrock) filter cubes, and a 40X 0.8 NA water immersion objective (LUMPLFLN, Olympus). Perforated-patch recordings were made using parafilm-wrapped borosilicate pipettes filled with internal solution containing 135 mM KCl, 10 mM NaCl, 2 mM MgCl_2_, 10 mM HEPES, 0.5 mM EGTA, 0.1 mM CaCl_2_, 280 mOsm, pH 7.4 adjusted with KOH. Recording electrodes (6-10 MΩ) were back filled with internal solution, then filled with internal containing 100 µg/mL gramicidin (Sigma). Pipette capacitance neutralization and bridge balance were manually adjusted prior to current clamp recordings. ChAT-Cre+ neurons expressing rsChRmine were optically stimulated with 10 ms TTL pulses every 10 s using orange (596 nm) light ranging from 0.2 to 3 mW delivered through the microscope objective by an LED driver (Thorlabs). Baseline axonal responses to rsChRmine stimulation in ACSF were measured for 3 minutes and the average response from the first 10 sweeps at baseline was compared to the average response from the final 10 sweeps during a 3 min period in the presence of 1 µM DHβE (Tocris). Acute application of DHβE was achieved via gravity-driven bath perfusion. Current clamp recordings were sampled at 20 kHz and filtered at 10 kHz using a MultiClamp 700B and Digidata 1550B and analyzed using pClamp11 (Molecular Devices, Sunnyvale, CA). Data are presented as mean value ± SEM, where n or N represent number of cells and animals, respectively.

### Immunohistochemistry

Mice were transcardially perfused with 4% (w/v) paraformaldehyde (PFA) in PBS and brains were removed and post-fixed overnight at 4°C. A vibratome (Leica Biosystems) was used to prepare 50 µm coronal sections. After three 10-minute washes in PBS on a shaker, the slices were incubated with blocking solution (10% normal goat serum, 0.2% bovine serum albumin, and 0.5% Triton X-100 in PBS) for 1 hour. After one 10-minute wash in PBS, slices were incubated in primary antibodies using a concentration of 1:1000 for 12-20 hours at 4°C on a shaker. Primary antibodies include: rat mCherry monoclonal antibody (Invitrogen, M11217), chicken anti-GFP (Aves labs, GFP-1020), mouse anti-tyrosine hydroxylase (Millipore, MAB318), and anti-2A peptide (Millipore, MABS2005). After three washes of 10 minutes in PBS, secondary antibodies were added at a concentration of 1:750 and incubated for 2 hours at room temperature on a shaker. Secondary antibodies include: goat anti-rat Alexa Fluor 594 (Invitrogen, A11007), goat anti-chicken Alexa Fluor 488 (Abcam, AB150169), and goat anti-mouse Alexa Fluor 647 (Invitrogen, A21235). Both primary and secondary antibodies were incubated with slices in a carrier solution (1% normal goat serum, 0.2% bovine serum albumin, and 0.5% Triton X-100 in PBS). After three more washes, the slices were mounted on SuperFrost Plus glass slides with DAPI Fluoromount-G mounting medium (Southern Biotech) for visualization using a Nikon A1 confocal microscope or a Keyence BZ-X800 microscope.

### Data analysis

Subject mice were excluded (<5%) from the analysis if they did not reach behavioral criteria or on the basis of histology if they had (1) inaccurate implant placement or (2) off-target or minimal transgene expression. MATLAB (MathWorks) scripts from Tucker-Davis Technologies were used for signal processing. Signals were down-sampled by a factor of 10, underwent LOESS smoothing (window size = 30 ms) to reduce high-frequency noise, and analyzed in 20 s windows around the following timestamps: the first nosepoke of each trial (excluding FR1 trials, where the first nosepoke triggers reward delivery), reward delivery (which is coincident with a light-sound cue), and reward consumption (defined as when the mice first entered the magazine port after sucrose reward delivery). Entire sessions were debleached according to a previously-published iterative method^52^ which calculates the dF/F in short moving windows, centers and normalizes these windows, and then repeats these calculations for 100 temporally-offset windows in the same session. Z-scores were then calculated using the mean and standard deviation of the signal spanning the entire session to minimize confounds in behavior contaminating the local baseline periods for trial-based methods of analysis. As a control, the analyses were all repeated in a different manner, using local baselines to calculate Z-scores, and the results were qualitatively identical.

To separate the contribution of individual behavioral events that occurred close in time, we (1) fit regression models to predict photometry data based on multiple variables (see below) and (2) examined neural activity on the subset of trials with large inter-trial intervals (e.g., >30 or 60 s), which revealed similar results (e.g., Extended Data Fig. 2, 6, and 7). All trials within a session were first combined into a single session average, and then sessions were averaged together for each mouse. All photometry figures in the manuscript show the mean and standard error of the photometry signal across mice, which is a more conservative approach than using the trial- or session-average. To quantify neural activity, we used area under the curve—calculated using the trapezoidal numerical integration of the Z-scores for the windows defined in the figure legends—or calculated the maximum value in the windows denoted in the figure legends. Windows for quantification were chosen based on visual inspection of the traces and applied consistently throughout the analysis for any direct comparison between traces. To minimize bleaching confounds, we removed the first 5 minutes of each recording, when the steepest bleaching was likely to occur. We also limited our interpretations to short windows of data, avoiding any analysis of longer-timescale changes, which are more likely to be confounded by bleaching or other gradual changes (e.g., slight adjustments in the connection between the implant and the patch cord), and more likely to vary depending on the exact debleaching strategy used. In addition, for GRAB DA recordings we took the approach reported by Sych et al.^53^ and examined simultaneously-recorded control signals (405 nm), which we found to be flat or slightly negative traces with substantially lower amplitude than what we observed with the experimental excitation (465 nm). Although this analysis is imperfect because 405 nm is not the isosbestic point for GRAB DA, the result implies that motion artifacts, bleaching, or other intrinsic, non-DA dependent signal changes could not have made a major contribution to our results.

### Kernel regression analysis

To quantify the contribution of behavioral variables to neural activity, linear mixed effects models were used as in prior work^13,54–57^. First, task-relevant behavioral predictors (e.g., active nosepoke, inactive nosepoke, reward delivery, magazine entry, fixed-ratio, inter-trial interval) were aligned to the photometry signal^13,54,58^. Behavioral events were assigned as fixed effects, represented in a predictor matrix X of dimension N x p, where N is the number of timestamps in the photometry recording and p is the number of predictor variables^13^. Variables were represented in binary form, indicating whether a behavioral event occurred at a given timestamp. The mouse from which the data was collected was treated as a random effect, stored in a sparse design matrix Z of size N x m, where m is the number of mice from which data was collected. The predictor matrix X was time shifted by T_1_ (T_1_=60 timeshifts/1.5 s, sucrose; T_1_=100 timeshifts/2.5 s, ChRmine) timestamps backwards and T_2_ (T_2_=125 timeshifts/4 s, sucrose; T_2_=400 timeshifts/10 s, ChRmine) timestamps forward to encapsulate effects of behavioral variables on the signal at previous and future timestamps^13^. The new, time-shifted predictor matrix was of dimension N x p(2(T_1 +_ T_2_) + 1). Data for timestamps at which no behavioral event of interest occurred, or which did not fall within the [-T_1_, T_2_] timestamps of a behavioral event, was excluded. The fixed effects matrix X and random effects matrix Z were used to train a linear mixed effects model of the form y = Xβ + Zµ + ε, where y is the N x 1 dimensional response vector consisting of the photometry signal values at each timestamp, β is a p(2(T_1 +_ T_2_) + 1) x 1 vector of beta coefficients for each time-shifted predictor variable, µ is an m x 1 vector of coefficients for each random effect, and ε is an N x 1 vector representing the residual variance that cannot be explained by the fixed or random effects. Response kernels were generated and plotted over the time window for each behavioral variable representing each behavioral variable’s contribution to the predicted fiber photometry signal^13,54,58^. The observed signal was plotted against the predicted signal to demonstrate the efficacy of the model fit. The model was trained using 10-fold cross validation; reported R^2^ values and plotted beta coefficients are the average across model runs for all 10-folds. The relative contribution of each behavioral variable to the total photometry signal was determined by performing a leave-one-out analysis where behavioral variables were sequentially removed from the model to observe the change in the models’ overall predictive ability as quantified by the R^2^ value. The relative contribution of a variable was then defined as the proportional change in R^2^ when that variable was removed from the model^55^.

### Quantification and statistical analysis

Investigators were blinded to the manipulations that experimental subjects had undergone during the behavioral testing, recordings and data analysis. All photometry analysis and behavioral analysis was conducted in MATLAB. Quantifications of the photometry and behavioral results were graphed and analyzed using GraphPad. All data were tested for normality of sample distributions using the Shapiro-Wilk normality test and when violated, non-parametric statistical tests were used. Normally distributed data were analyzed by paired, two-tailed t-tests, or one- or two-factor repeated measures ANOVA. Non-normally distributed data was analyzed using the Wilcoxon matched-pairs signed rank test, or Friedman’s test. When data were not paired, unpaired t-tests or the Mann-Whitney rank test was performed for normally and non-normally distributed data, respectively. If individual data points were missing from these matched comparisons, mixed-effects models were used instead. Mixed-effects models were also used when examining the effects of multiple fixed effects (e.g., FR and drug) accounting for the random effect of subject. In these cases, the significance of the fixed effects was reported in the figure legends and if there were significant fixed effects, Sidak-corrected post-hoc comparisons were reported with asterisks in the figures. Kruskal-Wallis tests were used to compare three or more unmatched groups (e.g., DA responses from trials with different inter-trial intervals). Spearman correlations were used to measure the association between two independently-measured observations (e.g., rewards on consecutive days). All statistical tests were two-sided and performed in MATLAB (Mathworks) or Prism (GraphPad). NS, not significant. *P < 0.05, **P < 0.01, ***P < 0.001, ****P < 0.0001. In all figures, data are shown as mean ± s.e.m.

## Acknowledgements

We thank the Eshel Lab, Malenka Lab, and Amitai Shenhav for critical discussion, and Paul Kramer for technical assistance with dopamine axon recordings. We thank Yulong Li for providing gACh4m virus. We also thank the Stanford Gene Vector and Virus core for reagents. This work was supported by philanthropic funds donated to the Nancy Pritzker Laboratory at Stanford University. G.C.T. was supported by the Stanford Medicine Berg Scholars Program. M.B.P. was supported by NIH grant K99 DA056573. Z.Z. was supported by the Stanford Wu Tsai Neurosciences Institute. R.C.M. was supported by a grant from the Stanford Wu Tsai Neurosciences Institute, a grant from the UCSF Dolby Family Center for Mood Disorders, and NIH grant P50 DA042012. N.E. was supported by NIH grant K08 MH123791, a Brain & Behavior Research Foundation Young Investigator Grant, a Burroughs Wellcome Fund Career Award for Medical Scientists, and a Simons Foundation Bridge to Independence Award.

## Author Contributions

Conceptualization: GCT, MBP, RCM, NE

Methodology: GCT, MBP, TY, VM, ND, ZZ, NE

Investigation: GCT, MBP, TY, VM, ND, ZZ

Visualization: MBP, GCT, NE

Funding acquisition: MBP, RCM, NE

Project administration: GCT, NE

Supervision: RCM, NE

Writing – original draft: MBP, GCT, NE

Writing – review & editing: GCT, MBP, TY, VM, ND, ZZ, RCM, NE

## Data availability

The datasets generated and analyzed in this study are available from the corresponding author upon reasonable request.

## Code availability

Code used for data processing and analysis is available from the corresponding author upon reasonable request.

## Correspondence

Correspondence and requests for materials should be addressed to Neir Eshel.

---

**Extended Data Fig 1.**
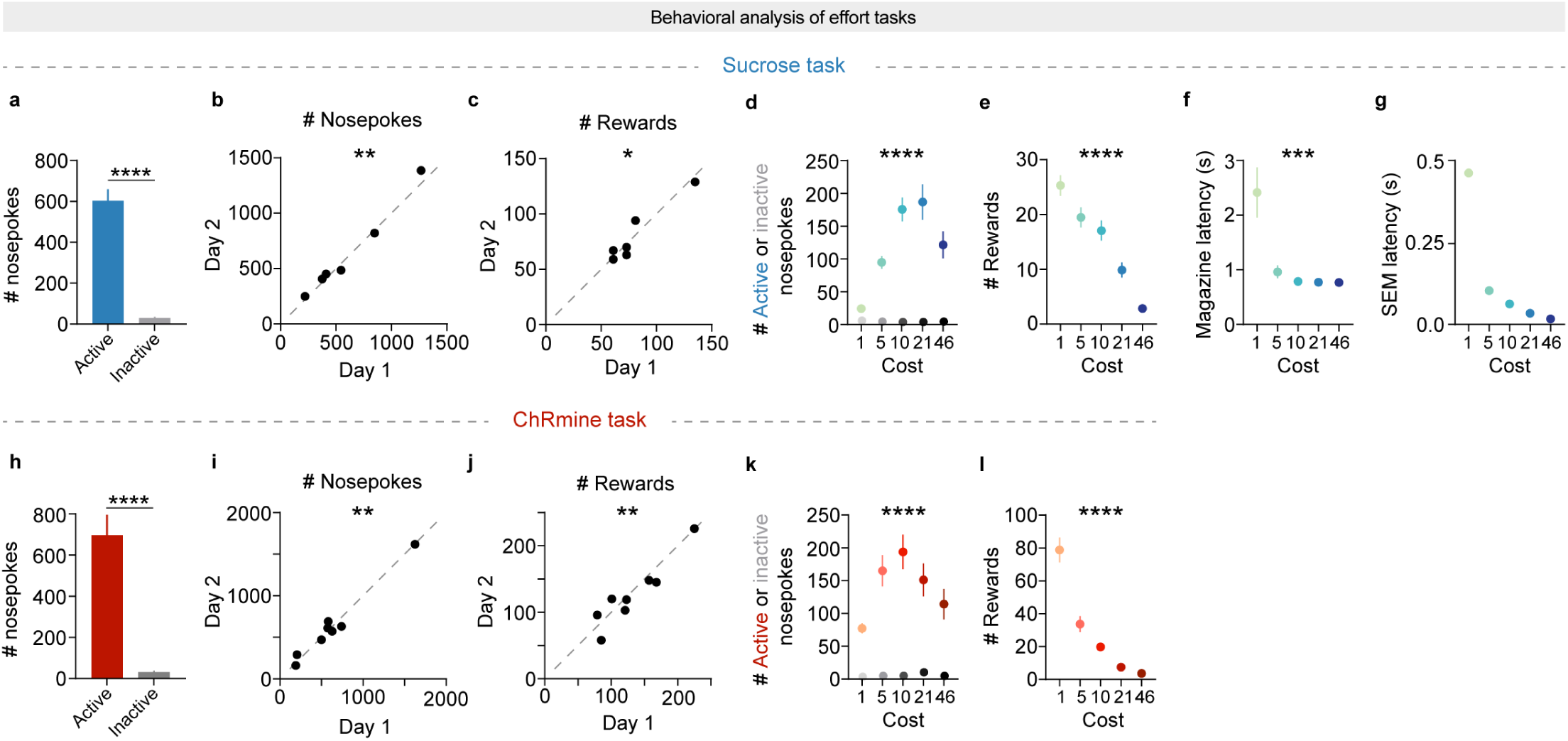
Mice readily perform the sucrose and ChRmine effort tasks. **a.** Number of active and inactive nosepokes in the sucrose effort task. Paired t-test, t = 9.98, df = 28, ****p < 0.0001, *n* = 6. **b.** Nosepoking consistency on the last two days of training. Spearman correlation, r = 1.00, **p = 0.0028, *n* = 6 (12 sessions from 6 mice). **c.** Reward earning consistency on the last two days of training. Spearman correlation, r = 0.85, *p = 0.044, *n* = 6 (12 sessions from 6 mice). **d.** Number of active and inactive nosepokes by FR in the sucrose effort task. Active: Friedman’s test, Friedman statistic = 53.23, ****p < 0.0001; n = 6. Inactive: Friedman’s test, Friedman statistic = 6.26, p = 0.18; *n* = 6. **e.** Number of rewards by FR in the sucrose effort task. Friedman’s test, Friedman statistic = 83.82, ****p < 0.0001; *n* = 6. **f.** Magazine latency by FR in the sucrose effort task. Mixed effects model, F(4,1540) = 4.80, ***p = 0.0008; *n* = 6. **g.** SEM of magazine latency by FR in the sucrose effort task. **h-l**. Similar to A-G except for the self-stimulation task. Statistics for **h**: Paired t-test, t = 6.73, df = 47, ****p < 0.0001, *n* = 8. Statistics for **i**: Spearman correlation, r = 0.88, **p = 0.0072, *n* = 8 (16 sessions from 8 mice). Statistics for **j**: Spearman correlation, r = 0.88, **p = 0.0072, *n* = 8 (16 sessions from 8 mice). Statistics for **k**: Active: Friedman’s test, Friedman statistic = 47.36, ****p < 0.0001; *n* = 8. Inactive: Friedman’s test, Friedman statistic = 8.55, p = 0.073; *n* = 8. Statistics for **l**: Friedman’s test, Friedman statistic = 168.1, ****p < 0.0001; *n* = 8.

**Extended Data Fig 2.**
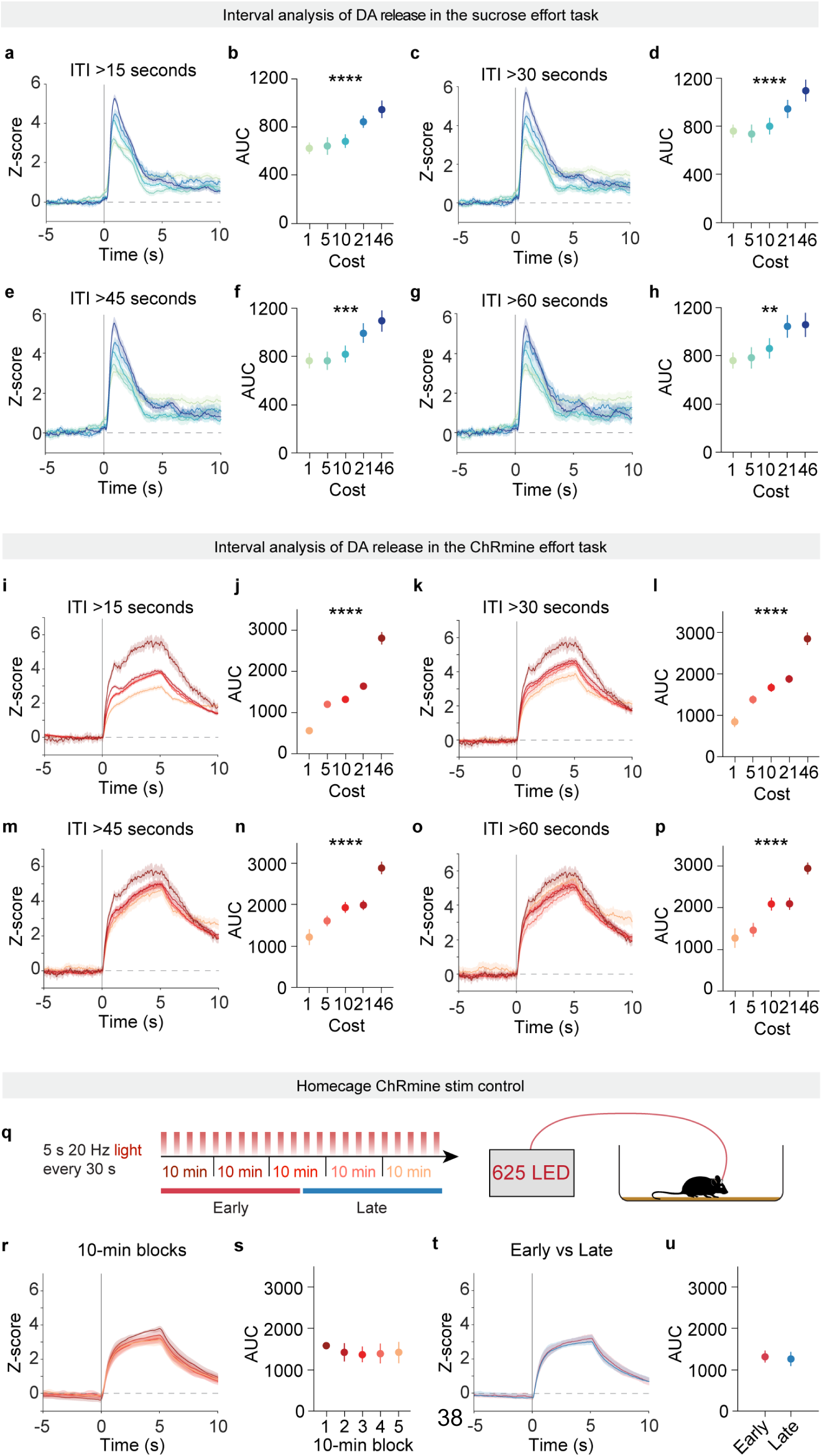
Inter-trial interval does not account for increased DA release after high effort expenditure. **a.** GRAB DA activity in the sucrose effort task for trials separated by an inter-trial interval (ITI) of >15 s. **b.** Area under the curve (0-4 s) for each FR for trials separated by ITI of >15 s. Mixed-effects model, main effect of FR, F(4,331) = 6.48, ****p < 0.0001, *n* = 6. **c.** GRAB DA activity in the sucrose effort task for trials separated by an ITI of >30 s. **d.** Area under the curve (0-4 s) for each FR for trials separated by ITI of >30 s. Mixed-effects model, main effect of FR, F(4,321) = 6.25, ****p < 0.0001, *n* = 6. **e.** GRAB DA activity in the sucrose effort task for trials separated by an ITI of >45 s. **f.** Area under the curve (0-4 s) for each FR for trials separated by ITI of >45 s. Mixed-effects model, main effect of FR, F(4,305) = 4.82, ***p = 0.0009, *n* = 6. **g.** GRAB DA activity in the sucrose effort task for trials separated by an ITI of >60 s. **h.** Area under the curve (0-4 s) for each FR for trials separated by ITI of >60 s. Mixed-effects model, main effect of FR, F(4,238) = 6.48, **p = 0.010, *n* = 6. **i-p**. Similar to **A-H** except for self-stimulation task. Statistics for **J**: Mixed-effects model, main effect of FR, F(4,802) = 60.36, ****p < 0.0001, *n* = 8. Statistics for **L**: Mixed-effects model, main effect of FR, F(4,388) = 47.22, ****p < 0.0001, *n* = 8. Statistics for **N**: Mixed-effects model, main effect of FR, F(4,231) = 18.85, ****p < 0.0001, *n* = 8. Statistics for **P**: Mixed-effects model, main effect of FR, F(4,153) = 14.01, ****p < 0.0001, *n* = 8. **q.** Schematic of the homecage ChRmine stimulation control. **r.** GRAB DA activity in the homecage stimulation control split by 10-minute blocks. **s.** Area under the curve (0-6 s) for each block. Friedman’s test, Friedman statistic = 4.20, p = 0.41; *n* = 4. **t.** GRAB DA activity in the homecage control split by first versus second-half stimulations. **u.** Area under the curve (0-6 s) for early versus late stimulations. Wilcoxon matched-pairs signed rank test, sum of signed ranks = -10.00, p = 0.13; *n* = 4.

**Extended Data Fig 3.**
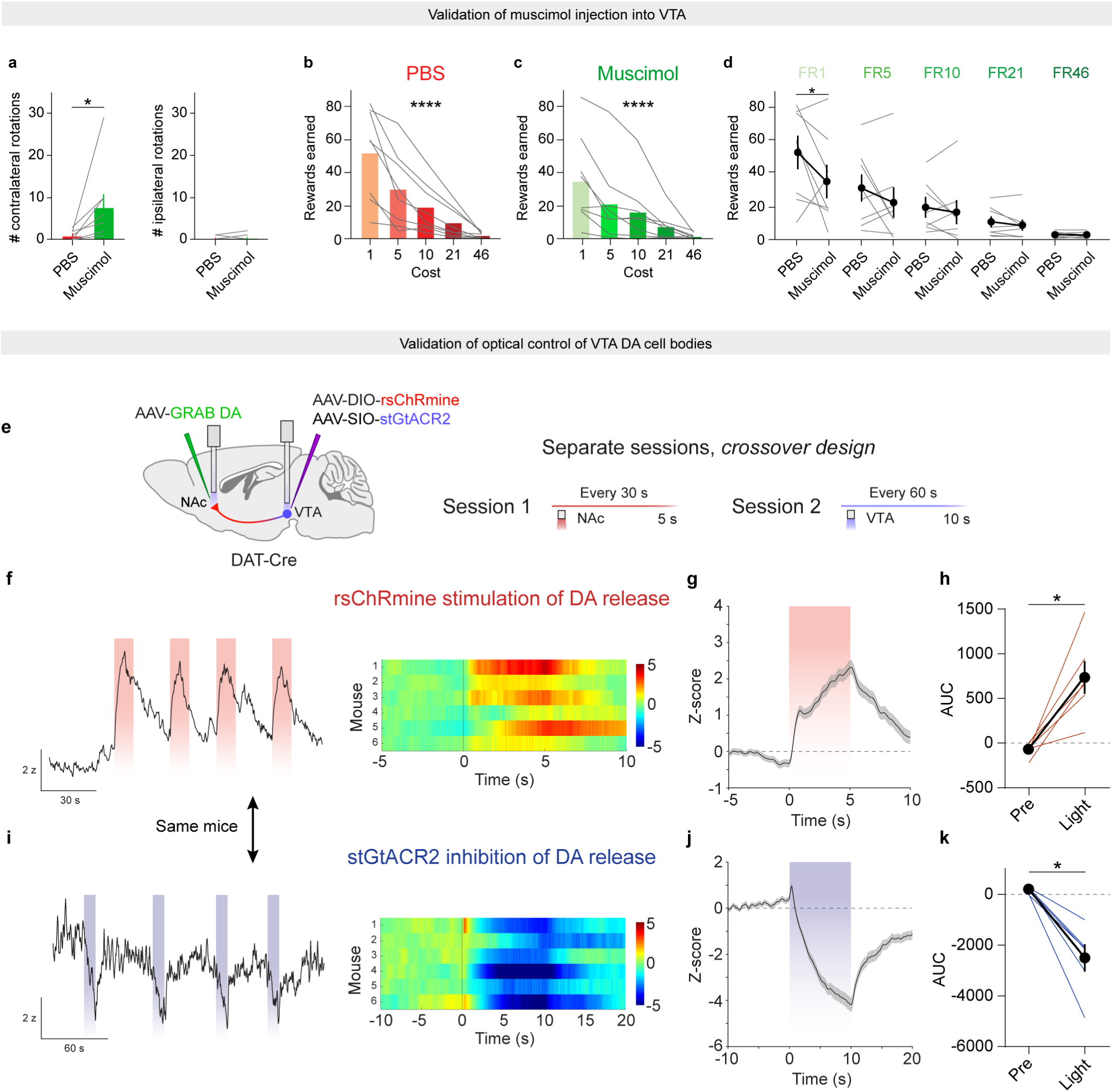
Validation of VTA DA cell body manipulations. **a.** Left, number of contralateral rotations mice made in the first 5 min of the ChRmine effort task when either PBS or muscimol was microinjected into the VTA. Wilcoxon matched-pairs signed rank test, sum of signed ranks = 28.00, *p = 0.016, *n* = 8. Right, number of ipsilateral rotations mice made in the same time. Wilcoxon matched-pairs signed rank test, sum of signed ranks = 2.00, p > 0.99, *n* = 8. **b.** Rewards earned for each FR after VTA microinjection with PBS. Friedman test, Friedman statistic = 31.12, ****p < 0.0001; *n* = 8. **c.** Rewards earned for each FR after VTA microinjection with muscimol. Friedman test, Friedman statistic = 25.83, ****p < 0.0001; *n* = 8. **d.** Rewards earned for each FR with pair-wise comparisons between PBS and muscimol. Mixed-effects model, fixed effect of FR, F(4,28) = 19.82, ****p < 0.0001; fixed effect of drug, F(1,7) = 1.86, p = 0.21; interaction of FR and drug; F(4,28) = 2.46; p = 0.068. Sidak-corrected multiple comparisons: FR1 PBS vs muscimol, *p = 0.037; FR5 PBS vs muscimol, p = 0.55; FR10 PBS vs muscimol, p > 0.99; FR21 PBS vs muscimol, p > 0.99; FR46 PBS vs muscimol, p > 0.99; *n* = 8. **e.** Schematic of AAV injection, fiber implantation, and experimental design for the validation of both ChRmine stimulation and stGtACR2 inhibition. **f.** Left, representative trace of GRAB DA activity with ChRmine stimulation. Right, heatmap of GRAB DA activity with ChRmine stimulation. **g.** GRAB DA signal during ChRmine stimulation averaged across all mice (*n* = 6). **h.** Area under the curve of GRAB DA signal before (-5-0 s) and during (0-5 s) ChRmine stimulation. Wilcoxon matched-pairs signed rank test, sum of signed ranks = 21.00, *p = 0.031, *n* = 6. **i.** Left, representative trace of GRAB DA activity with stGtACR2 inhibition. Right, heatmap of GRAB DA activity with stGtACR2 inhibition. **j.** GRAB DA signal during stGtACR2 inhibition averaged across all mice (*n* = 6). **k.** Area under the curve of GRAB DA signal before (-10-0 s) and during (0-10 s) stGtACR2 inhibition. Wilcoxon matched-pairs signed rank test, sum of signed ranks = -21.00, *p = 0.031, *n* = 6.

**Extended Data Fig 4.**
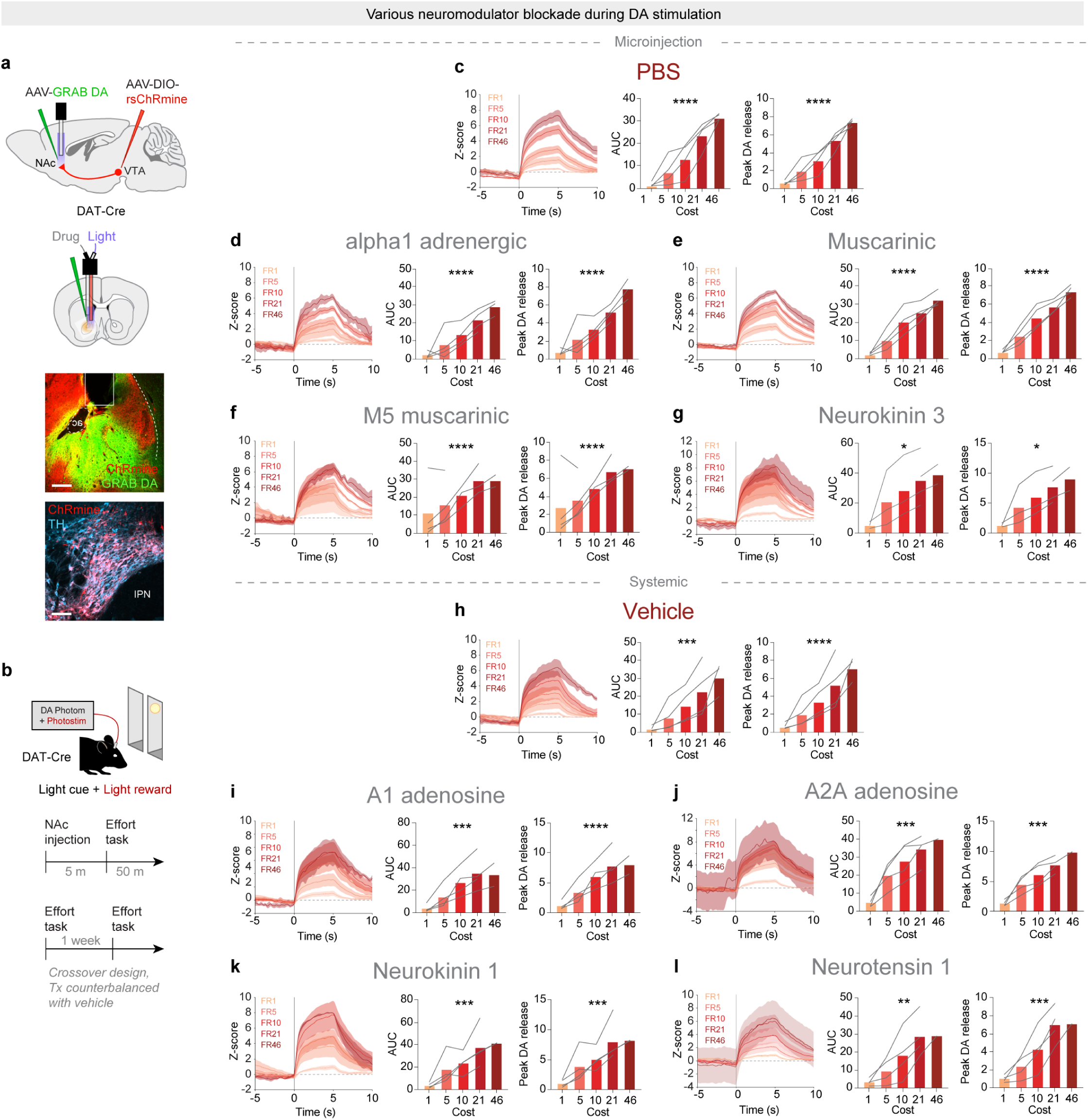
Effects of neuromodulator blockade on DA encoding of effort. **a.** Top, schematic of AAV injection and fiber-cannula implant for the self-stimulation task with injection of receptor antagonist. Bottom, representative images of NAc fiber placement and viral expression of ChRmine. **b.** Schematic of effort task and experimental schedules. **c-g**. Average GRAB DA activity, area under the curve (AUC, 0-6 s), and peak DA release for each FR during the self-stimulation task with microinjection of PBS (**c**, AUC: Friedman’s test, Friedman statistic = 16.00, ****p < 0.0001; *n* = 4; Peak: Friedman’s test, Friedman statistic = 16.00, ****p < 0.0001; *n* = 4), terazosin (**d**, AUC: Mixed effects model, F(4,11) = 21.96, ****p < 0.0001; *n* = 4; Peak: Mixed effects model, F(4,11) = 34.15, ****p < 0.0001; *n* = 4), scopolamine (**e**, AUC: Friedman’s test, Friedman statistic = 16.00, ****p < 0.0001; *n* = 4; peak: Friedman’s test, Friedman statistic = 16.00, ****p < 0.0001; *n* = 4), ML375 (**f**, AUC: Mixed effects model, F(4,8) = 34.06, ****p < 0.0001; *n* = 4; peak: One-way ANOVA, F(4,8) = 30.32, ****p < 0.0001; *n* = 4), and osanetant (**g**, AUC: Mixed effects model, F(4,7) = 5.28, *p = 0.028; *n* = 4; peak: Mixed effects model, F(4,7) = 7.18, *p = 0.013; *n* = 4). **h-l**. Average GRAB DA activity, AUC (0-6 s), and peak DA release for each FR during the self-stimulation task with systemic injection of vehicle corn oil/saline (**h**, AUC: Mixed effects model, F(4,11) = 12.92, ***p = 0.0004; *n* = 4; peak: Mixed effects model, F(4,11) = 18.29, ****p < 0.0001; *n* = 4), DPCPX (**i**, AUC: Mixed effects model, F(4,10) = 16.03, ***p = 0.0002; *n* = 4; peak: Mixed effects model, F(4,10) = 24.40, ****p < 0.0001; *n* = 4), istradeffyline (**j**, AUC: One-way ANOVA, F(4,12) = 10.01, ***p = 0.0008; *n* = 4; peak: One-way ANOVA, F(4,12) = 15.20, ***p = 0.0001; *n* = 4), aprepitant (**k**, AUC: Mixed effects model, F(4,10) = 14.16, ***p = 0.0004; *n* = 4; peak: Mixed effects model, F(4,10) = 16.52, ***p = 0.0002; *n* = 4), and SR 48692 (**l**, AUC: Mixed effects model, F(3,9) = 13.21, **p = 0.0012; *n* = 4; peak: Mixed effects model, F(3,9) = 20.79, ***p = 0.0002; *n* = 4).

**Extended Data Fig 5.**
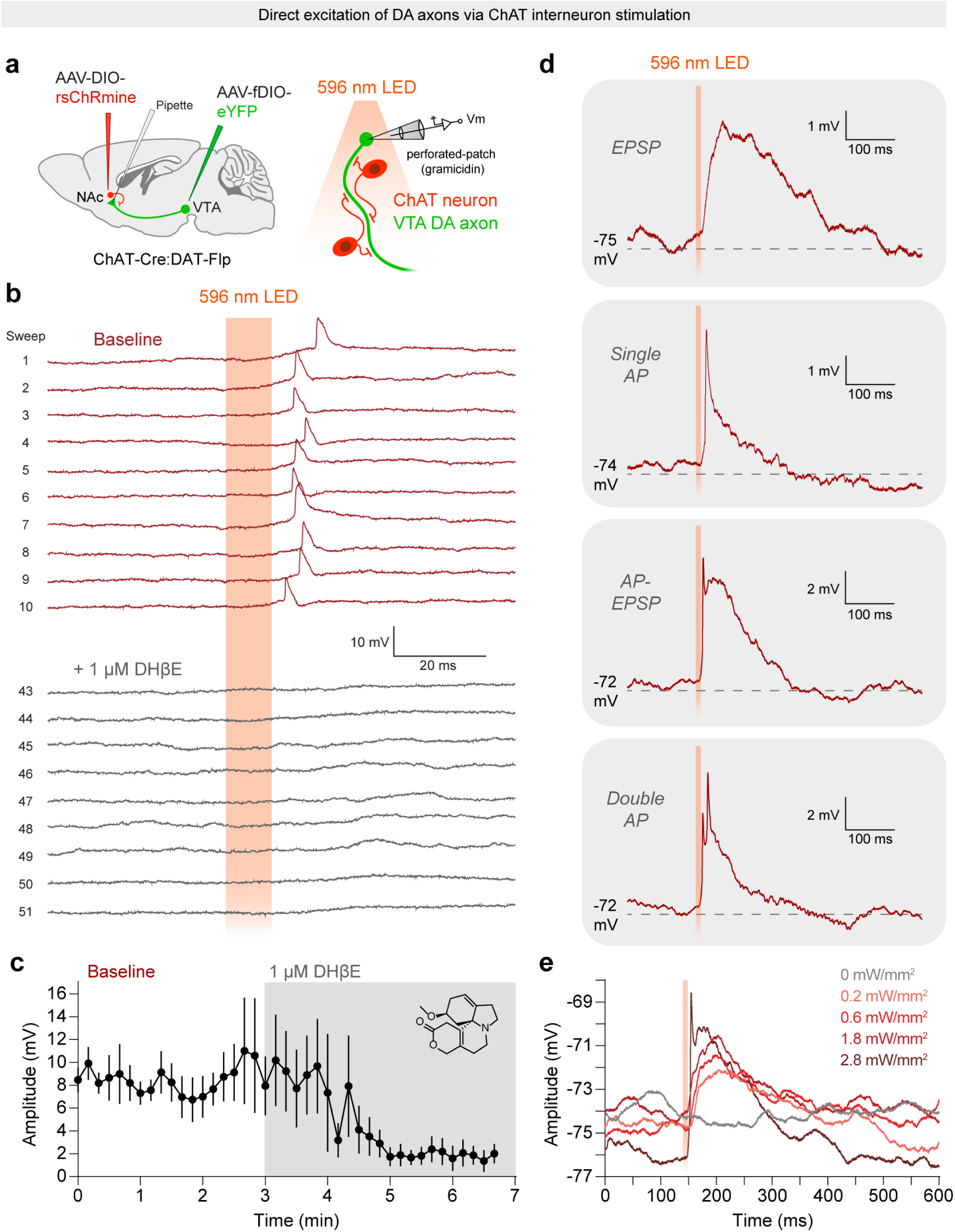
Additional electrophysiology data from DA axon recordings. **a.** Schematic of AAV injection and electrophysiological recording configuration. **b.** Representative sweeps from a single recording of an eYFP+ VTA DA axon bleb in response to rsChRmine stimulation of cholinergic interneurons at baseline and in the presence of 1 µM DHβE. **c.** Time course of average axonal action potential (axAP) amplitude evoked by rsChRmine stimulation before and during acute perfusion of 1 µM DHβE; *n* = 3 axons, N = 3 mice. **d.** Representative traces showing VTA DA axon responses to rsChRmine stimulation of cholinergic interneurons, including axEPSPs, axAPs, compound axAP-EPSPs, and axAP bursts. **e.** Representative DA axon bleb recording showing a relationship between the irradiance of cholinergic rsChRmine stimulation and axonal response amplitude.

**Extended Data Fig 6.**
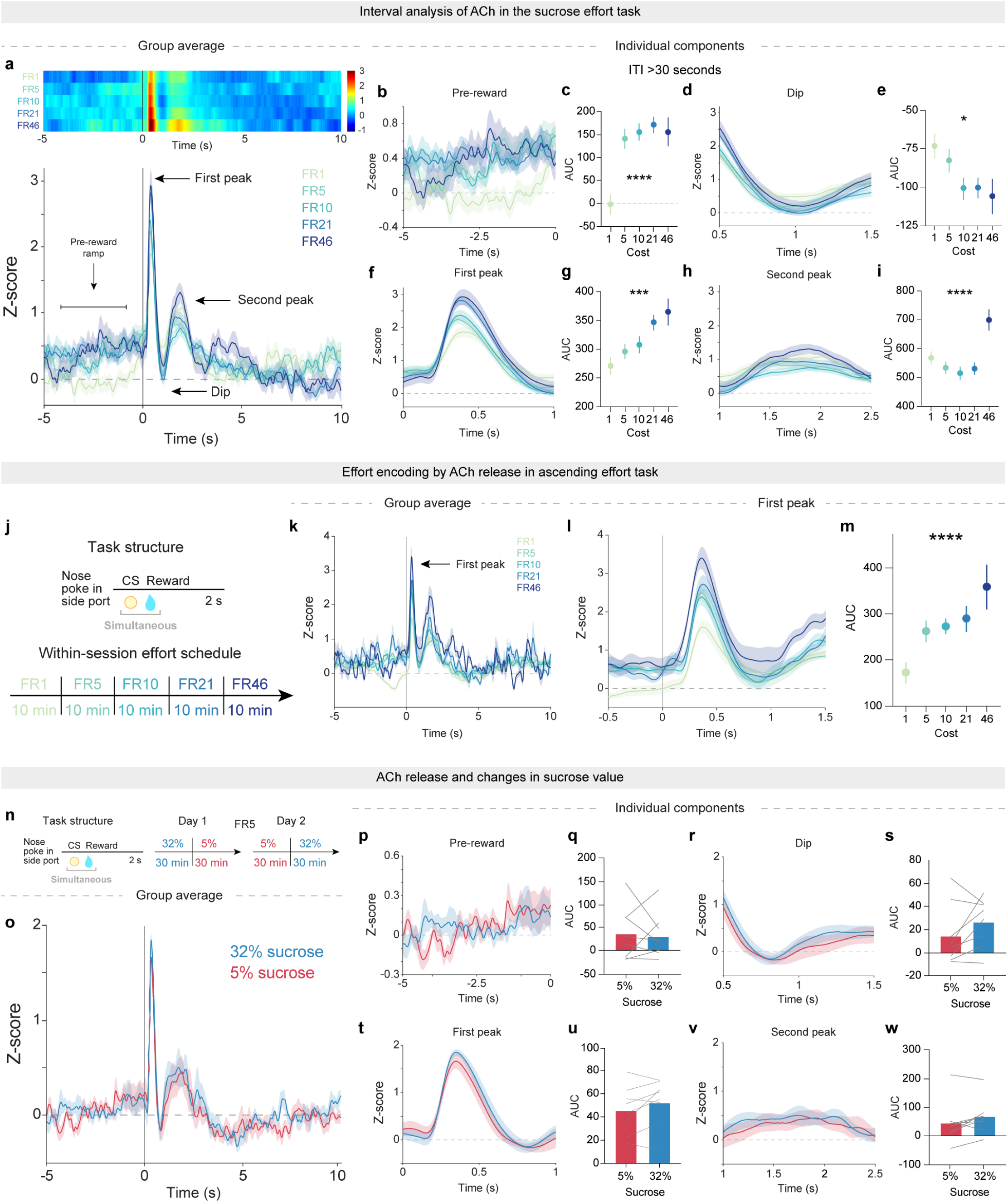
Inter-trial interval and task structure do not account for increased ACh release after high effort expenditure. **a.** Top, heatmap and bottom, averaged traces of GRAB ACh activity during the sucrose effort task indicating the pre-reward ramp, first peak, dip, and second peak (*n* = 8). **b.** GRAB ACh activity during the pre-reward ramp for trials separated by >30 s. **c.** Area under the curve (AUC, -5 to 0 s) for each FR during the pre-reward period. One-way ANOVA, F(4,497) = 9.68, ****p < 0.0001; *n* = 8. **d.** GRAB ACh activity during the dip for trials separated by >30 s. **e.** AUC (0.75-1.25 s) during the dip. One-way ANOVA, F(4,497) = 2.74, *p = 0.028; *n* = 8. **f.** GRAB ACh activity during the first peak for trials separated by >30 s. **g.** AUC (0.25-0.75 s) for each FR during the first peak. One-way ANOVA, F(4,497) = 5.39, ***p = 0.0003; *n* = 8. **h.** GRAB ACh activity during the second peak for trials separated by >30 s. **i.** AUC (1-2.5 s) for each FR during the second peak. One-way ANOVA, F(4,497) = 7.21, ****p < 0.0001; *n* = 8. **j.** Task design for the ascending FR sucrose effort task. **k.** GRAB ACh activity during the ascending sucrose effort task indicating the first peak. **l.** Closer view of the first peak of GRAB ACh activity during the ascending effort task. **m.** AUC (0.25-0.75 s) for each FR during the first peak. Mixed effects model, F(4,23) = 9.90, ****p < 0.0001; *n* = 8. **n.** Task design for changing the concentration of sucrose reward in an FR5 task. **o.** GRAB ACh activity at reward delivery for 5% and 32% sucrose earned at FR5 (*n* = 8). **p.** GRAB ACh activity during the pre-reward ramp for 5% and 32% sucrose rewards. **q.** AUC (-5-0 s) for each FR pre-reward. One-way ANOVA, F(4,497) = 9.68; p < 0.0001. Wilcoxon matched-pairs signed rank test, sum of signed ranks = -2.00, p = 0.95; *n* = 8. **r.** GRAB ACh activity during the dip for 5% and 32% sucrose rewards. **s.** AUC (0.75-1.25 s) for each FR during the dip. Wilcoxon matched-pairs signed rank test, sum of signed ranks = 20.00, p = 0.20; *n* = 8. **t.** GRAB ACh activity during the first peak for 5% and 32% sucrose rewards. **u.** AUC (0.25-0.75 s) for each FR during the first peak. Wilcoxon matched-pairs signed rank test, sum of signed ranks = 24.00, p = 0.11; *n* = 8. **v.** GRAB ACh activity during the second peak for 5% and 32% sucrose rewards. **w.** AUC (1-2.5 s) for each FR during the second peak. Wilcoxon matched-pairs signed rank test, sum of signed ranks = 22.00, p = 0.15; *n* = 8.

**Extended Data Fig 7.**
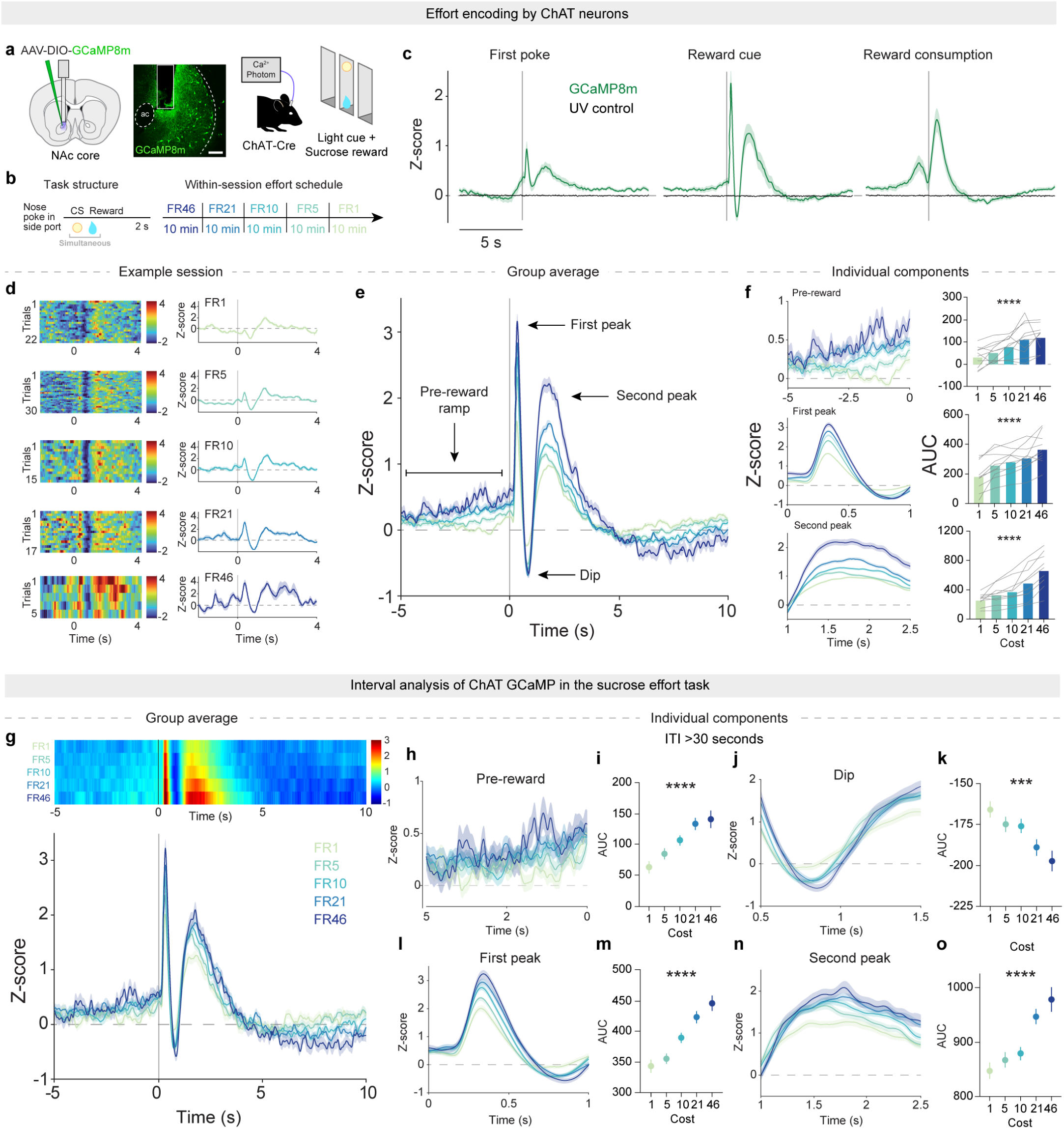
NAc cholinergic interneuron activity encodes effort and closely resembles ACh release. **a.** Schematic of AAV injection and representative image for the recording of ChAT GCaMP activity in the sucrose effort task. **b.** Task structure for recording ChAT GCaMP activity during the sucrose effort task. **c.** ChAT GCaMP activity aligned to the first nosepoke, reward delivery, and reward consumption (*n* = 10 mice). **d.** ChAT GCaMP heat maps and average traces aligned to reward delivery from a single example session. **e.** ChAT GCaMP activity averaged across all mice and sessions during the sucrose effort task indicating the pre-reward ramp, first peak, dip, and second peak. **f.** ChAT GCaMP activity during the pre-reward, first peak and second peak of ChAT GCaMP response to reward delivery. Area under the curve for each FR during the pre-reward (-5-0 s) period, Friedman test, Friedman statistic = 24.16, ****p < 0.0001; during first peak (0.25-0.75 s), Friedman test, Friedman statistic = 28.48, ****p < 0.0001; and during second peak (1-2.5 s), Friedman test, Friedman statistic = 37.36, ****p < 0.0001; *n* = 10. **g.** Top, heatmap and bottom, averaged traces of ChAT GCaMP activity across all mice (*n* = 10) during the sucrose effort task, restricted to trials with inter-trial interval (ITI) > 30 s. **h.** ChAT GCaMP activity during the pre-reward period of the reward delivery response for trials with ITI >30 s. **i.** Area under the curve of the ChAT GCaMP activity during the pre-reward period (-5-0 s) for trials with ITI >30 s. One-way ANOVA, F(4,1199) = 10.89, ****p < 0.0001, *n* = 10. **j.** GRAB ACh activity during the dip for trials with ITI > 30 s. **k.** Area under the curve of the GRAB ACh activity during the dip (0.75-1.25 s) for trials with ITI >30 s. One-way ANOVA, F(4,1199) = 5.23, ***p = 0.0004, *n* = 10. **l.** ChAT GCaMP activity during the first peak of the reward delivery response for trials with ITI >30 s. **m.** Area under the curve of the ChAT GCaMP activity during the first peak (0.25-0.75 s) for trials with ITI >30 s. One-way ANOVA, F(4,1199) = 19.45, ****p < 0.0001, *n* = 10. **n.** ChAT GCaMP activity during the second peak of the reward delivery response for trials with ITI > 30 s. **o.** Area under the curve of the ChAT GCaMP activity during the second peak (1-2.5 s) for trials with ITI >30 s. One-way ANOVA, F(4,1199) = 12.50, ****p < 0.0001, *n* = 10.

**Extended Data Fig 8.**
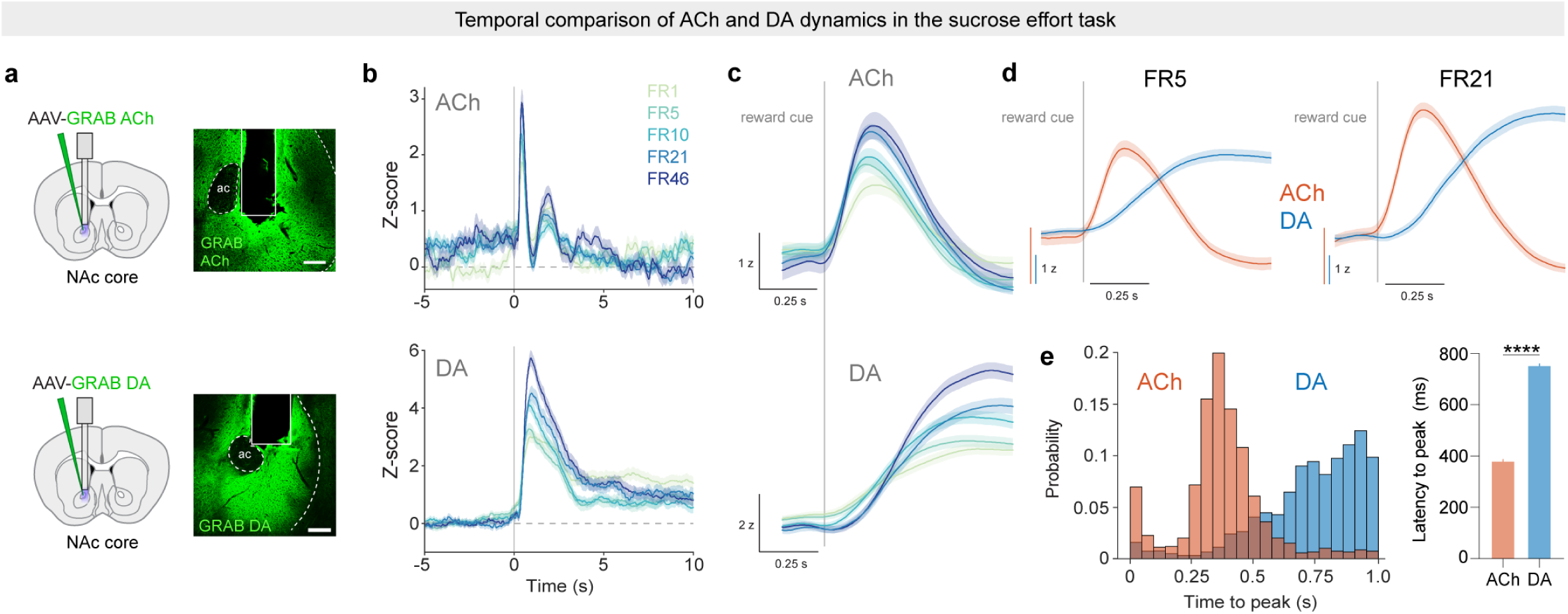
Temporal comparison of ACh and DA. **a.** Schematic of AAV injections and representative images for the sucrose effort task with recording of GRAB ACh (top) and GRAB DA (bottom) activity. Scale bar = 100µm. **b.** GRAB ACh (top) and GRAB DA (bottom) activity averaged across all mice and sessions during the sucrose effort task. Replotted from Figs. 1E and 4E. **c.** Enlarged plot of the GRAB ACh (top) and GRAB DA (bottom) activity averaged across all mice and sessions during the sucrose effort task depicting the early period of the reward delivery response. **d.** Enlarged plot of the FR5 and FR21 GRAB ACh and DA activity overlayed to depict the early reward delivery response period. **e.** Left, histogram of the latency to peak for all trials in the GRAB ACh and GRAB DA recordings during the sucrose effort task. Right, bar graph comparing the average latency to peak for GRAB ACh and GRAB DA during the sucrose effort task. Unpaired t-test, t = 58.18, df = 4225, ****p < 0.0001, *n* = 14.

**Extended Data Fig 9.**
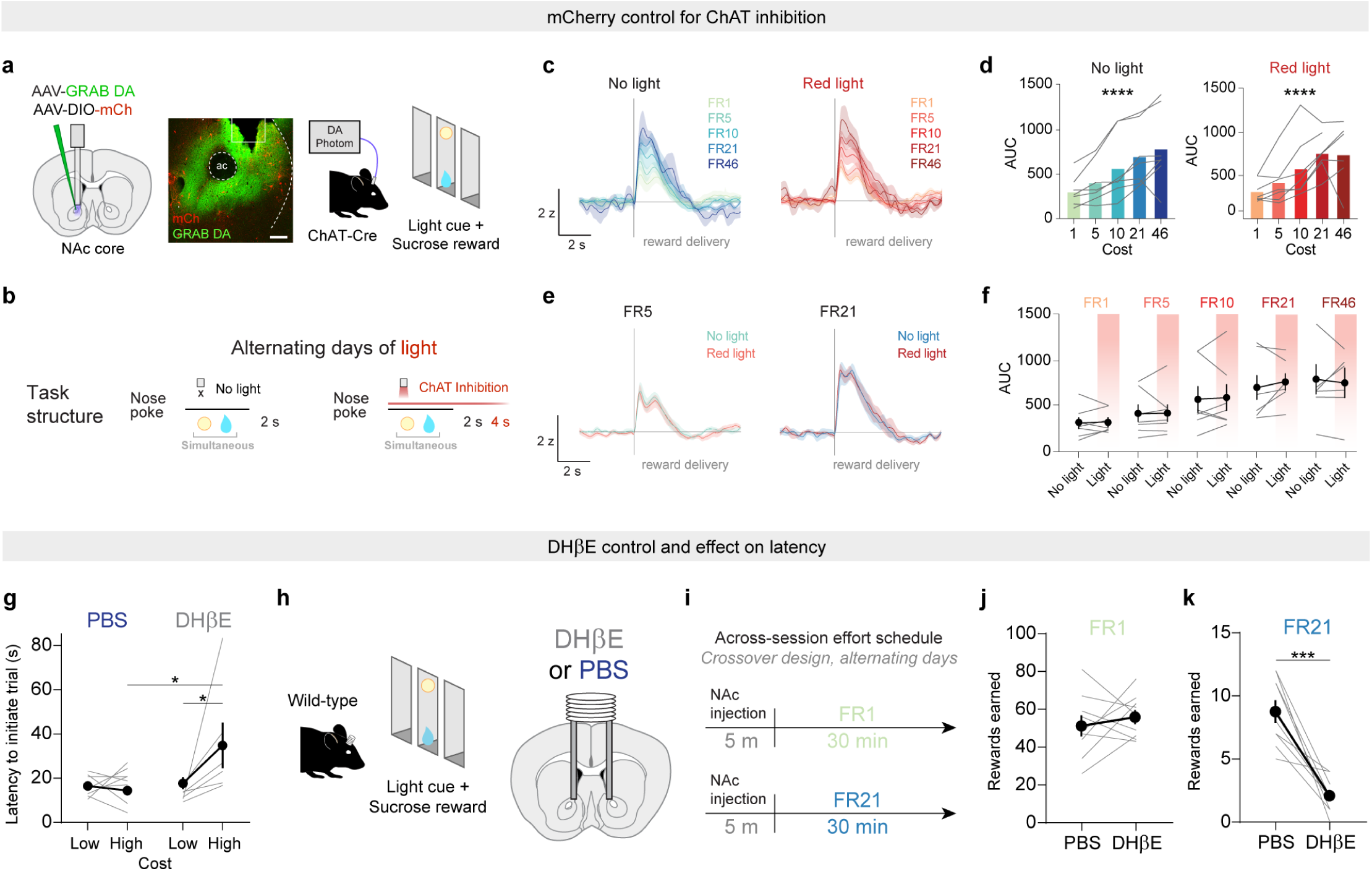
Behavioral controls for ACh manipulations. **a.** Schematic of AAV injections and representative image for mCherry red light control of cholinergic interneurons with GRAB DA recording. Scale bar = 100µm. **b.** Task design for red light control of cholinergic neuron inhibition sucrose effort task. **c.** GRAB DA activity during the sucrose task during no light and red-light sessions (*n* = 7). **d.** Area under the curve (0-4 s) for each FR in no light sessions and red-light sessions. No light, Friedman test, Friedman statistic = 20.46, ****p < 0.0001; *n* = 7. Red light: mixed-effects model, F(4,23) = 9.90, ****p < 0.0001; *n* = 7. **e.** GRAB DA activity during the sucrose effort task for no light and red-light sessions comparing FR5 and FR21. **f.** Area under the curve (0-4 s) for each FR comparing no light to red light. Mixed-effects model, fixed effect of FR, F(4,24) = 14.55, ****p < 0.0001; fixed effect of red-light, F(1,6) = 0.36, p = 0.57; fixed effect of FR and red-light interaction; F(4,23) = 0.23; p = 0.92. Sidak-corrected multiple comparisons: FR1 no light vs red-light, p = 0.99; FR5 no light vs red-light, p > 0.99; FR10 no light vs red-light, p = 0.99; FR21 no light vs red-light, p = 0.80; FR46 no light vs red-light, p = 0.99; *n* = 7. **g.** Latency to initiate the next trial after reward delivery with PBS vs DHβE microinjection during the sucrose effort task, comparing low effort (FR1 and FR5) vs high effort (FR10, 21, and 46) trials. Mixed-effects model, fixed effect of effort, F(1,26) = 2.69, p = 0.11; fixed effect of drug, F(1,26) = 5.54, *p = 0.026; interaction of effort and drug; F(1,26) = 4.37; *p = 0.047. Sidak-corrected multiple comparisons: low vs high effort PBS, p = 0.93, low vs high effort DHβE, *p = 0.034; *n* = 9. **h.** Schematic of cannula implant for the sucrose control task with PBS or DHβE microinjection. **i.** Task design for sucrose control task in which mice were injected with PBS or DHβE before performing an entire session at either FR1 or FR21. **j.** Rewards earned after PBS vs DHβE microinjection in FR1 sessions. Wilcoxon matched-pairs signed rank test, W = 11.00, p = 0.57; *n* = 9. **k.** Rewards earned after PBS vs DHβE microinjection in FR21 sessions. Wilcoxon matched-pairs signed rank test, W = -45.00, ***p = 0.0039; *n* = 9.

